# Chromatin Interaction Neural Network (ChINN): A machine learning-based method for predicting chromatin interactions from DNA sequences

**DOI:** 10.1101/2020.12.30.424817

**Authors:** Fan Cao, Yu Zhang, Yichao Cai, Sambhavi Animesh, Ying Zhang, Semih Akincilar, Yan Ping Loh, Wee Joo Chng, Vinay Tergaonkar, Chee Keong Kwoh, Melissa J. Fullwood

## Abstract

Chromatin interactions play important roles in regulating gene expression. However, the availability of genome-wide chromatin interaction data is limited. Various computational methods have been developed to predict chromatin interactions. Most of these methods rely on large collections of ChIP-Seq/RNA-Seq/DNase-Seq datasets and predict only enhancer-promoter interactions. Some of the ‘state-of-the-art’ methods have poor experimental designs, leading to over-exaggerated performances and misleading conclusions. Here we developed a computational method, Chromatin Interaction Neural Network (ChINN), to predict chromatin interactions between open chromatin regions by using only DNA sequences of the interacting open chromatin regions. ChINN is able to predict CTCF-, RNA polymerase II- and HiC-associated chromatin interactions between open chromatin regions. ChINN also shows good across-sample performances and captures various sequence features that are predictive of chromatin interactions. To apply our results to clinical patient data, we applied CHINN to predict chromatin interactions in 6 chronic lymphocytic leukemia (CLL) patient samples and a cohort of open chromatin data from 84 CLL samples that was previously published. Our results demonstrated extensive heterogeneity in chromatin interactions in patient samples, and one of the sources of this heterogeneity were the different subtypes of CLL.

## Introduction

Chromatin interactions play important roles in regulating gene expression^1, 2^. They bridge enhancers to genes^3-5^ and create insulated domains to constrain the reach of enhancers^6^. High-throughput experimental techniques such as high-throughput Chromosome Conformation Capture (Hi-C)^7^ and Chromatin Interaction Analysis with Paired-End Tags (ChIA-PET)^8^ have been developed to detect genome-wide chromatin interactions. These techniques greatly advanced the understanding of genome organization and its roles in transcription regulation^3, 9-11^. However, due to costs and technical challenges, these methods have not been widely applied to large cohorts of cell lines or clinical samples. Hence, our understanding of how common or rare chromatin interactions are in different patient samples is limited.

A predictor that uses DNA sequences to predict chromatin interactions could potentially expand our understanding of genome organization. Sophisticated computational methods such as DeepSea^12^ and DeepBind^13^ have demonstrated that many transcription factor binding sites in open chromatin regions could be predicted from DNA sequences. Additionally, various computational methods have been developed to predict chromatin interactions to complement the experimental techniques^14-20^. Many of these methods rely on using various functional genomics data including chromatin immunoprecipitation sequencing (ChIP-seq) data of transcription factors and histone modifications, open chromatin data, and transcription data^14, 16, 18, 20^. Methods such as RIPPLE^16^, TargetFinder^18^, and JEME^14^ reported high performances in predicting enhancer-promoter interactions using supervised machine learning approaches. Although the reported performances were exaggerated by using cross-validation with random splitting of samples^21^, these methods suggested that chromatin interactions could be potentially predicted from 1-dimensional functional genomics data^22^.

Recently, the convolutional neural network framework was adapted to predict Hi-C contact matrix from 1-dimentional sequence data in a method called “Akita” ^23^. CTCF-associated genome folding pattern can be observed in the prediction results, suggesting the importance of CTCF in regulating chromatin interactions. In addition, prediction results can recapture the differences in genome folding between a normal and genetically altered cell lines, indicating that machine learning framework can predict different genome folding profiles given different input DNA sequences.

However, there are several limitations. First, Akita only performs predictions with 1 Mb DNA sequence regions, thus long-range chromatin interactions cannot be predicted. Second, it is unclear whether ChIA-PET data can be predicted. Third, this method was not tested for its ability to predict chromatin interactions *de novo* in patient cancer samples. In this study, we investigated the possibility of utilizing DNA sequence features to predict chromatin interactions between open chromatin regions, regardless of distance between them. We demonstrated that open chromatin interactions can be predicted accurately from functional genomic data at the resolutions of the experimental techniques. We then developed a novel method, called Chromatin Interaction Neural Network (ChINN) to predict open chromatin interactions from DNA sequences. This model has been developed for RNA Polymerase II ChIA-PET interactions, CTCF ChIA-PET interactions and Hi-C interactions. ChINN was able to identify convergent CTCF motifs, AP-1 transcription family member motifs such as FOS, and other transcription factors such as MYC as being important in predicting chromatin interactions.

Moreover, we further applied our model to a set of 6 newly generated chronic lymphocytic leukemia samples, which showed patient-specific chromatin interactions. We were able to validate predicted interactions by Hi-C. The models were then applied to a cohort of previously published 84 chronic lymphocytic leukemia (CLL) samples^24^. We found additional evidence for patient-specific chromatin interactions, and chromatin interactions that were different in different subtypes of CLL. Taken together, our results indicate that ChINN can predict chromatin interactions, and application of ChINN to cancer patient samples demonstrates widespread patient heterogeneity in chromatin interactions.

## Results

### Open chromatin interactions can be predicted from functional genomic features

In light of Xi et al.^21^ and our previous study^22^ showing that the existing prediction methods have exaggerated performances, we first tried to demonstrate that chromatin interactions could be predicted from functional genomic data. Many previous studies focused on enhancer-promoter interactions that were annotated using chromatin interactions derived from HiC or ChIA-PET ^14, 16, 18^. The enhancers used were typically hundreds of base pairs, while the chromatin interaction anchors were much larger in size. The resolution discrepancy could lead to the introduction of a lot of noises to the training datasets (Figure 1a). Thus, we used the chromatin interaction anchors directly.

**Figure 1:**
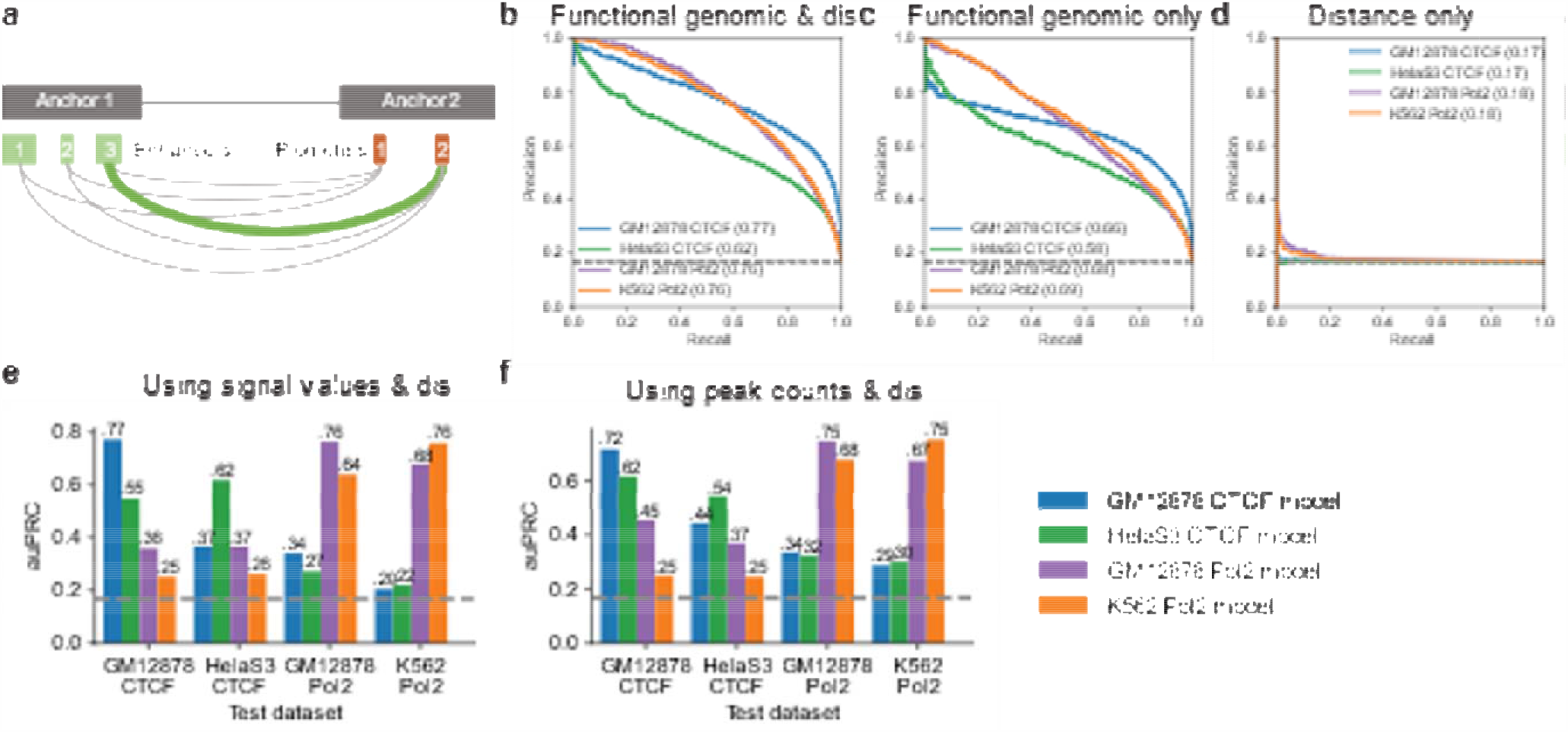
Performances of the functional genomic models on distance-matched datasets. **a**, Illustration of resolution discrepancy between cis-regulatory elements and chromatin interaction anchors. **b-d**, Precision-recall curves of the functional genomic models on distance-matched datasets using features based on **b**, functional genomic data and distance (dis); **c**, only functional genomic data; and **d**, only distance. Numbers in brackets indicate the area-under precision-recall curve. **e-f**, Across-sample performances using distance (dis) and **e**, signal values and **f**, peak counts.

Positive samples were constructed from ChIA-PET datasets separately and the corresponding distance-matched negative datasets were generated (Supplementary Figure 1). The resulting distance-matched datasets have positive-to-negative ratios of approximately 1:5 and all chromatin interactions were between open chromatin regions in the corresponding cell types. We used ChIP-seq data of transcription factors and histone modifications commonly available to GM12878, K562 and HelaS3 and DNase-seq data from ENCODE^25^ to annotate the anchors and build the feature vectors (Supplementary Table 1). For each chromatin interaction, the average signal of each transcription factor, histone modification and open chromatin were calculated for both anchors. The distance between two anchors was also used as a feature.

Gradient boosted trees^26^ were used to build models for each dataset. We tested three feature sets: 1) all common functional genomics data and distance; 2) distance only; and 3) common functional genomics data only. The models trained on all features achieved area under precision-recall curve (auPRC) ranging from 0.62 to 0.77 (Figure 1b), while models trained on distance are mostly at baseline (Figure 1d), showing that distance is properly controlled between positive and negative samples. The models trained on functional genomics features achieved auPRCs ranging from 0.58 to 0.69 (Figure 1c), lower than models trained on all features. These results showed that although distance alone cannot predict chromatin interactions, the interaction between distance and other features could help to distinguish between positive and negative chromatin interactions.

The across sample performances were lower than within-sample performances (Figure 1e). Using peak counts instead of signal values produced better across-sample performances but lower within-sample performances (Figure 1f). Models trained on RNA Polymerase II (Pol2) datasets generalize well to each other. Models trained on CTCF ChIA-PET datasets, however, did not generalize well to each other. Models trained on CTCF ChIA-PET data perform poorly on Pol2 ChIA-PET datasets and vice versa.

### Open chromatin interactions can be predicted from DNA sequences

Motivated by the results above, we went on to explore whether open chromatin interactions can be predicted from DNA sequences. We built a convolutional neural network, ChINN, to predict chromatin interactions between open chromatin regions using DNA sequences (Figure 2a). The models were trained on GM12878 CTCF, GM12878 Pol2, HelaS3 CTCF, K562 Pol2, and MCF-7 Pol2 datasets separately.

**Figure 2:**
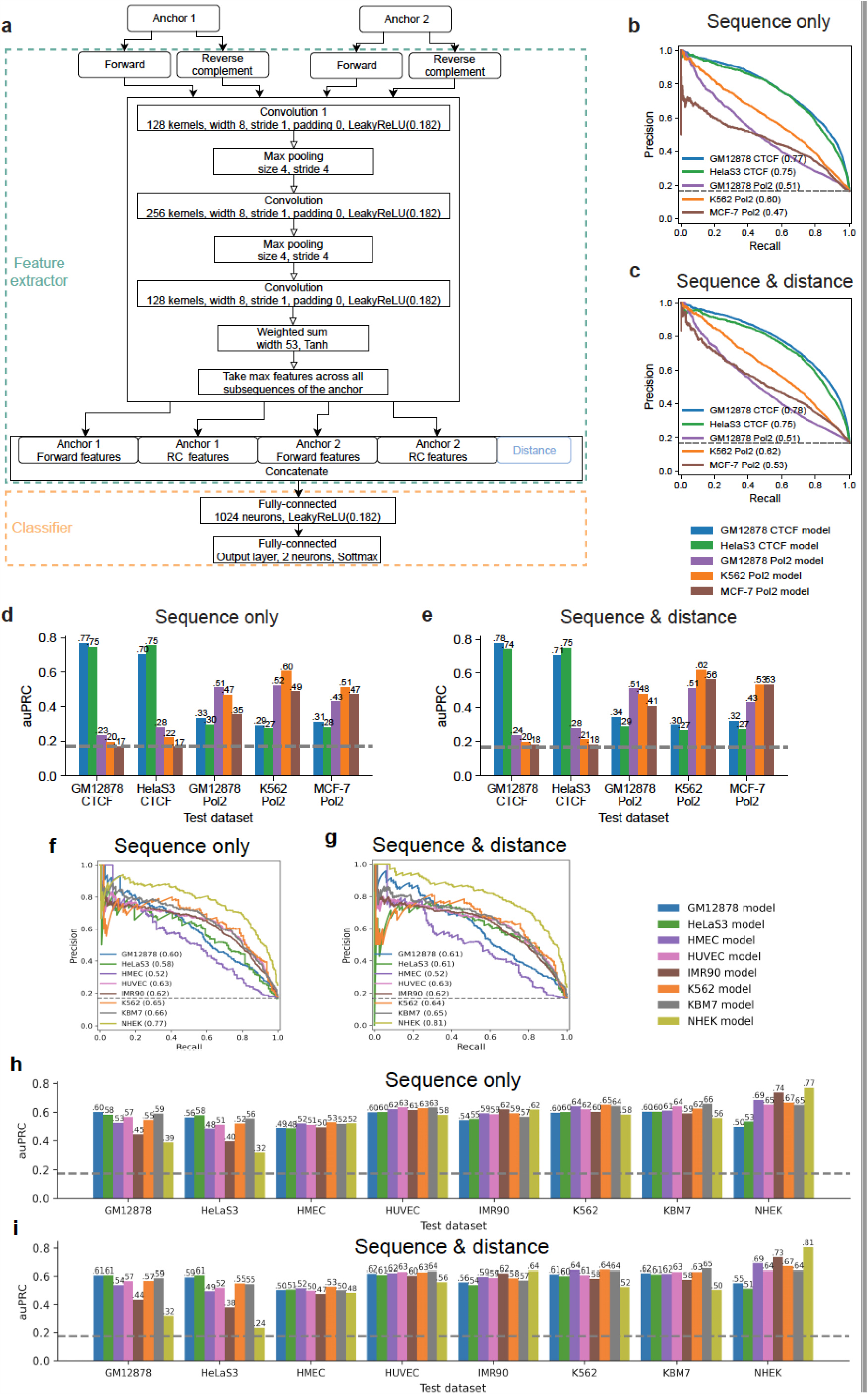
Architecture and performances of the sequence-based models on distance-matched datasets. **a**, The architecture of the sequence-based models using to train on distance-matched datasets. **b-c**, Precision-recall curves of the sequence-based models on distance-matched datasets using **b**, only sequence features or **c**, sequence features with distance. The numbers in the brackets indicates the area under precision-recall curves. **d-e**, Across-sample performances as measured by area-under precision-recall curve (auPRC) of the models on distance-matched datasets using **d**, only sequence features or **e**, sequence features with distance. **f-g**, Precision-recall curves of the sequence-based models on distance-matched HiC datasets using **f**, only sequence features or **g**, sequence features with distance. The numbers in the brackets indicates the area under precision-recall curves. **h-i**, Across-sample performances as measured by area-under precision-recall curve (auPRC) of the models on distance-matched HiC datasets using **h**, only sequence features or **i**, sequence features with distance.

Compared to using functional genomics data for prediction, using sequences produced better within-sample performances for CTCF ChIA-PET datasets with auPRCs of 0.77 for GM12878 CTCF and 0.75 for HelaS3 CTCF (Figure 2b), but worse within-sample performances for Pol2 ChIA-PET datasets with auPRC of 0.51 for GM12878 Pol2, 0.6 for K562 Pol2, and 0.47 for MCF-7 Pol2. Including distance as a feature to classifier only slightly improved the performances for the distance-matched datasets (Figure 2c). The across-sample performances of CTCF models showed well generalizability to each other (Figure 2d). Pol2 models can also generalize to each other. Models trained on CTCF ChIA-PET datasets perform poorly on Pol2 ChIA-PET datasets and vice versa (Figure 2d-e). The inability to generalize between CTCF chromatin interactions and Pol2 chromatin interactions could be attributed to the different sequence contexts.

For each model, we obtained and matched the position-weight matrices for all kernels on the first convolutional layer to known transcription factor binding motifs (Supplementary Figure 2). As expected, CTCF motif was captured by both CTCF models (Supplementary Figure 2a-b). Other than the CTCF motif, the remaining known transcription factor binding motifs learned by the two models were different. The patterns learned by Pol2 models showed more diversity and no matching transcription factor binding motif was shared among the three models (Supplementary Figure 2c-e). Interestingly, some of the transcription factors identified, such as ZNF143 in K562 and GATA3 in MCF-7, play important roles in the relevant cancer types^27, 28^.

Besides, we also trained CHINN model on GM12878, HeLaS3, HMEC, HUVEC, IMR90, K562, KBM7, and NHEK HiC data, respectively. The auPRCs of within-sample performances using only sequences range from 0.52 to 0.77 for the above eight HiC models (Figure 2f). Including distance as a feature to classifier only slightly improved the performances for the GM12878, HeLaS3, and NHEK HiC models (Figure 2g). The across-sample performances of all eight HiC models showed well generalizability to each other (Figure 2h-i).

Similarly, we obtained and matched the position-weight matrices for all kernels on the first convolutional layer to known transcription factor binding motifs for eight HiC datasets (Supplementary Table 2) and counted how many times each motif was detected (Supplementary Table 3). The CTCF motif was captured by all HiC models. The known transcription factor binding motifs learned by different HiC models were different. Some motifs, such as FOS, were learned by all models, but other motifs showed diversity, for example, ZN436 is detected by all other models except for HMEC, and ZIC3 is only detected by HeLaS3 (Supplementary Table 3).

### Convergent CTCF motifs are important for prediction of CTCF-associated open chromatin interactions

After extracting the sequence features from both the forward and reverse-complement sequences of the anchors, the sequence features were fed into the classifier to obtain a probability score that indicated how likely the pair of anchors were involved in a chromatin interaction. We obtained the feature importance scores of the gradient boosted trees trained and validated using a set of extended datasets that includes more negative samples than the distance-matched datasets (Methods, Supplementary Figure 3a-d). Distance was the most important feature in all models.

Next, we focused on the sequence features that were important for the prediction. Interestingly, in CTCF models the important sequence features were on different strands of the two anchors (Figure 3a-b), while Pol2 models did not show such pattern (Figure 3c-e). For the CTCF models, importance scores of features on different strands of the two anchors showed good correlation, while importance scores of features on the same strand of the two anchors did not show much correlation (Figure 3f). In contrast, the importance scores of features of Pol2 models were generally highly correlated regardless of the strand. The kernels on the last convolutional layer that generated the most important features in the extended CTCF models captured the CTCF motif (Supplementary Figure 3e-f), suggesting that convergent CTCF motifs were important for the prediction of CTCF-associated chromatin interactions. However, using only CTCF motif information for the prediction of CTCF-associated open chromatin interactions could not recapitulate the performance achieved by the convolutional neural network (Supplementary Figure 3g), indicating that CTCF was not the sole determining factor of chromatin interactions.

**Figure 3:**
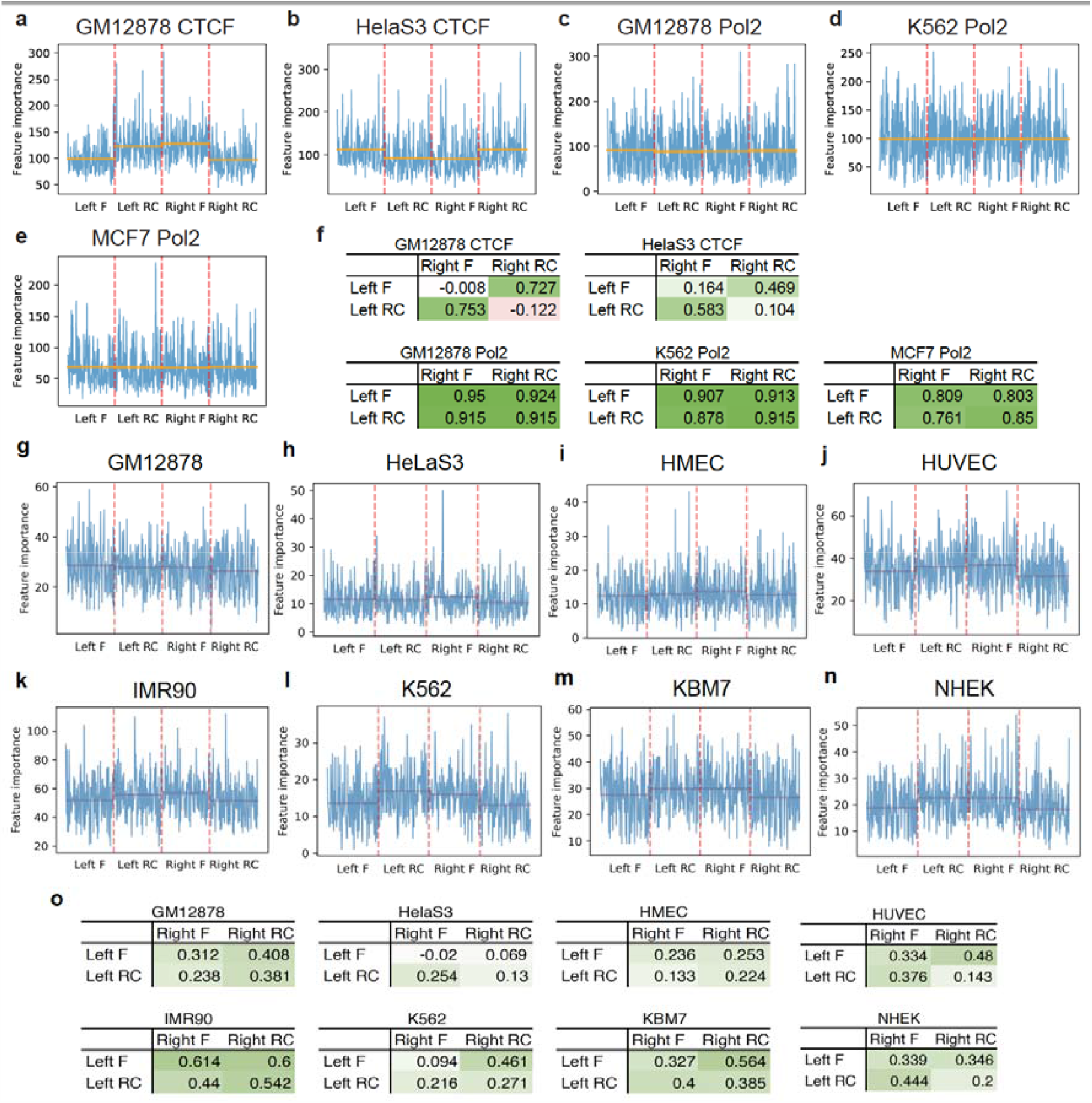
Sequence feature importance scores of gradient boosted trees trained on extended datasets. **a-e**, The importance scores of sequence features extracted from both directions (F: forward; RC: reverse complement) of the two anchors (left and right) by models trained on different datasets. The orange horizontal lines indicate average importance scores of the features from the strand of the anchor. **f**, Pearson correlations between feature importance scores of the two anchors. **g-n**, The importance scores of sequence features extracted from both directions (F: forward; RC: reverse complement) of the two anchors (left and right) by models trained on HiC datasets. The orange horizontal lines indicate average importance scores of the features from the strand of the anchor. **o**, Pearson correlations between feature importance scores of the two anchors in HiC datasets.

Similarly, we trained gradient boosted trees with the corresponding extended datasets for eight HiC datasets. Distance was still the most important feature in all models (Supplementary Figure 4a-d). When we visualized the sequence feature importance, although not as obvious as that of the CTCF models, we observed that the important sequence features were on different strands of the two anchors according to the corresponding mean values (Figure 3g-n). However, the importance scores of features did not show highly correlation on HiC datasets (Figure 3o). All the extended HiC models captured the CTCF motif via the kernels of the most important feature on the last convolutional layer (Supplementary Figure 4e), indicating that convergent CTCF motifs were important for the prediction of HiC data chromatin interactions.

### Predicting chromatin interactions from open chromatin regions

The above models were trained and evaluated on known chromatin interactions. Without knowledge of chromatin interactions, as is the case for many clinical samples and cell types, the locations of the anchors would not be known. To be able to predict chromatin interactions between open chromatin regions, the models need to be able to predict chromatin interactions between anchors constructed from open chromatin regions.

We tested different combinations of merging distances and extension sizes (Figure 4a) based on validation datasets and determined that the merging distance of 3000 bp and extension size of 1000 bp for the construction of anchors in GM12878 cells (Supplementary Figure 5a).

**Figure 4:**
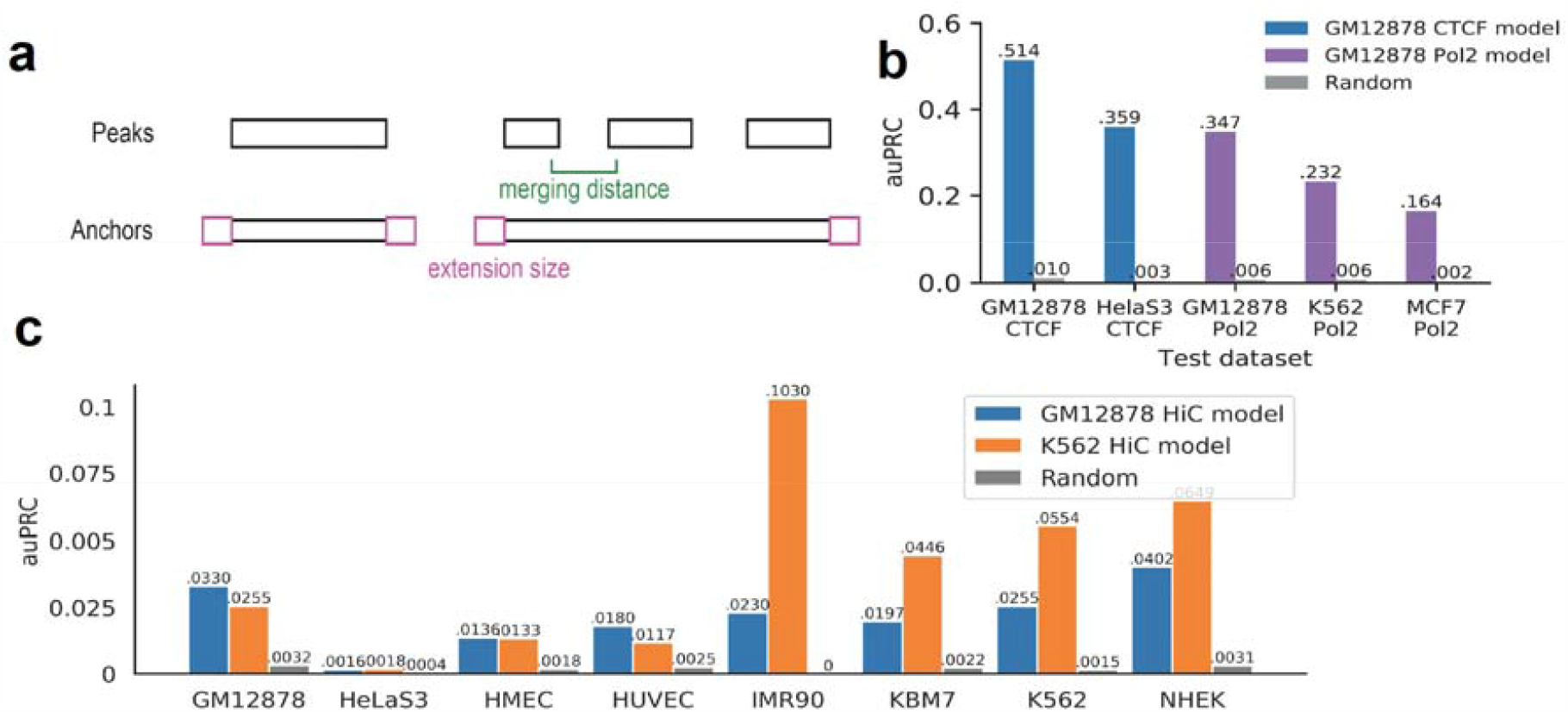
Performances of the from-DNase models and validations. **a**, Illustration of the two parameters, merging distance and extension size, used in constructing putative chromatin interactions anchors from open chromatin regions. **b**, Area under precision-recall curves of the from-DNase models. **c**, Area under precision-recall curves of the HiC from-DNase models.

The pairs generated between anchors constructed from open chromatin regions in GM12878 were used to train gradient boosted trees for both CTCF and Pol2 models (see Methods). The positive-to-negative ratios were about 1:122 for CTCF chromatin interaction labeled samples and 1:186 for Pol2 chromatin interaction labeled samples. The CTCF model achieved within-sample auPRC of 0.514 and the Pol2 model achieved auPRC of 0.347 (Figure 4b). In cross-sample evaluation, the CTCF model achieved auPRC of 0.359 on HelaS3 dataset and the Pol2 model achieved auPRCs of 0.232 and 0.164 on K562 and MCF-7 datasets, respectively (Figure 4b). We were able to validate some of the predicted chromatin interactions in MCF-7 cells using 4C-seq (Supplementary Figure 5b-d). Some of the validated chromatin interactions were not captured by the MCF-7 Pol2 ChIA-PET dataset, thus ChINN is able to identify bona fide chromatin interactions that might have been previously missed out due to insufficient sequence coverage.

We also generated pairs between anchors constructed from open chromatin regions in GM12878 and K562 HiC datasets with different combination of merging distance and extension size (Supplementary Figure 6a). We kept to use the same parameters as CTCF model, i.e. merging size of 3,000 and extension size of 1,000, to train gradient boosted trees due to the not significantly difference of auROC achieved by different parameters. The GM12878 and K562 HiC model achieved a little bit low auPRC in the within-sample and cross-sample evaluation (Figure 4c). However, the performances were acceptable when compared to the random auPRC values. Moreover, some of the predicted chromatin interactions in MCF-7 cells using 4C-seq were able to be validated by our HiC models (Supplementary Figure 6b-d).

### Exploring chromatin interactions in patient samples

Next, we wished to apply our machine learning methods to patient samples to understand if our method could predict chromatin interactions in a completely new dataset. We obtained 6 Chronic Lymphocytic Leukemia (CLL) patient samples. The clinical characteristics are described in Supplementary Table 4.

We prepared integrated Hi-C, ATAC-Seq and RNA-Seq libraries from these 6 samples. We used Juicer to call Topologically-Associated Domains and loops from these patient samples. Our CLL samples showed many TADs and loops (Supplementary Table 5), thus indicating that we were able to perform Hi-C in these patient samples.

Next, we applied GM12878 and K562 HiC models to six new CLL samples. The auPRC achieved by GM12878 HiC model range from 0.2772 to 0.4362, which are a bit higher than that of K562 HiC model, whose auPRC range from 0.2607 to 0.3996 (Figure 5a). We calculated the F-score with different thresholds and finally determined the threshold of 0.025 for GM12878 model and 0.016 for K562 model to make the prediction on new CLL samples (Supplementary Figure 7a-b), where the corresponding confusion matrix was shown as Figure 5b-c.

**Figure 5:**
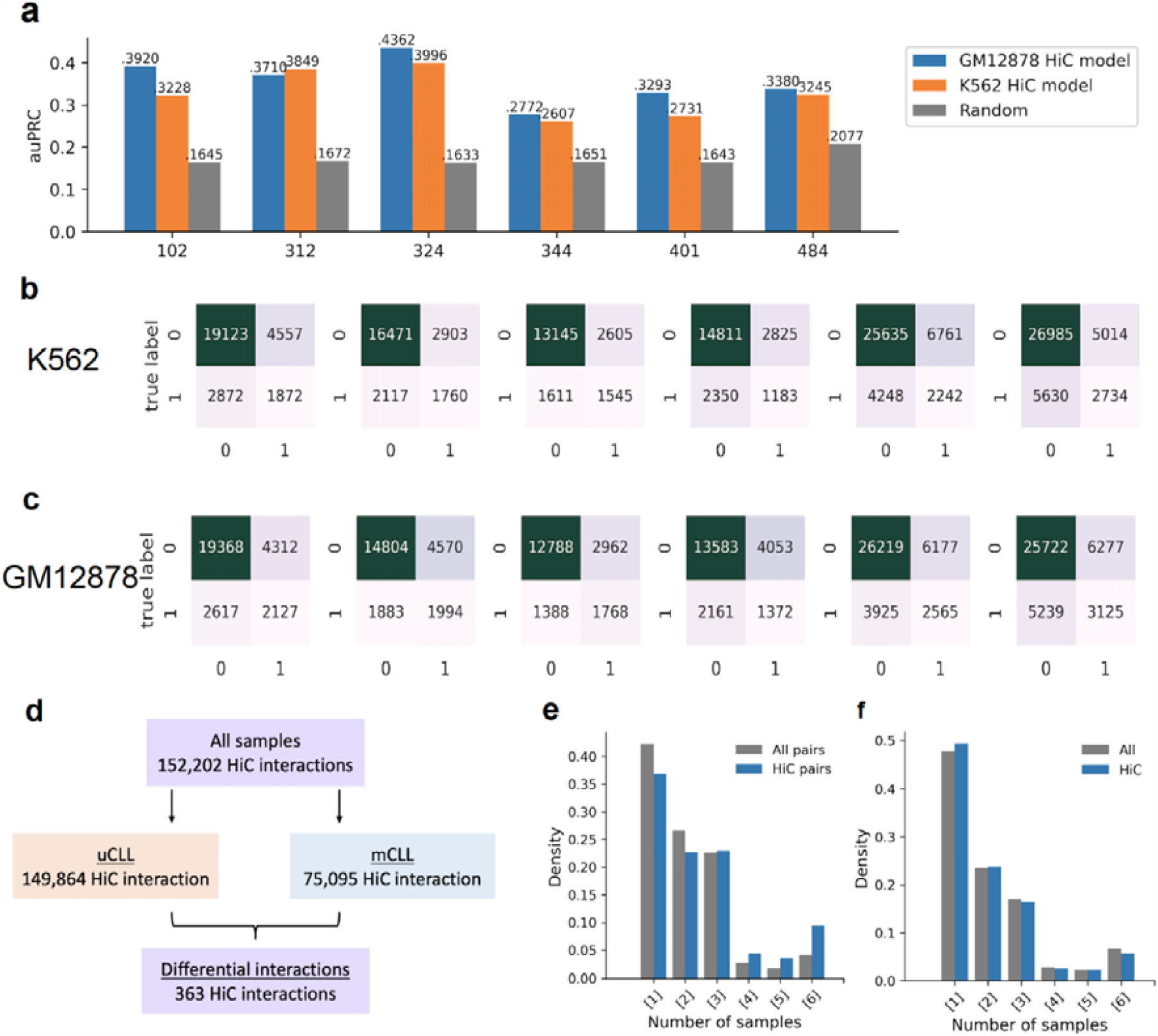
Applying HiC model on new CLL samples. **a**, the auPRC values achieved by GM12878 and K562 HiC model, x-axis: new CLL samples. **b-c**, The confusion matrices for 6 new CLL samples using K562 HiC model with threshold of 0.016 and GM12878 HiC model with threshold of 0.025. x-axis: true label, y-axis: predicted label. 0: negative, 1: positive. **d**, Summary of the predicted chromatin interactions in the 6 new CLL samples and the differential chromatin interactions between uCLL and mCLL samples. **e**, Conservation analysis of predicted chromatin interactions in new CLL samples. All pairs: all possible pairs used for prediction. **f**, Uniqueness analysis of open chromatin regions that overlap with HiC peaks from GM12878 cells in new CLL samples. All: all open chromatin regions.

With the selected threshold, a total of 152,202 HiC-associated open chromatin interactions were predicted (Figure 5d) by GM12878 HiC model. One question we asked was whether there is patient heterogeneity in Hi-C data. We found extensive patient heterogeneity (Figure 5e-f), as observed from the lack of conservation of chromatin interactions across the new CLL samples and the overlapping peaks between new CLL samples and GM12878 HiC peaks.

In addition, we also applied our ChINN framework on the six new CLL samples and built models using Hi-C and ATAC-seq data from each CLL sample. Our Hi-C libraries identified 1795 chromatin interactions unique in uCLL samples and 10663 chromatin interactions unique in mCLL samples (Figure 6a). Uniqueness analysis of the Hi-C interactions from these six CLL samples showed high patient heterogeneity (Figure 6b). These models have auPRC range from 0.37 to 0.58 (Figure 6c). In addition, across-sample testing of these CLL models on other datasets from other CLL sample suggest a comparable performance (Figure 6d). Inclusion of distance did not result in dramatic increase of the model performance (Supplementary Figure 8a-8b). Similarly, the first convolutional layers of all CLL models were able to capture the CTCF and AP-1 transcription family member (FOS, JUN, JUNB, JUND) binding motif (Supplementary Figure 8c) as the Hi-C models we showed earlier (Supplementary Figure 4e; Supplementary Table 2-3).

**Figure 6.**
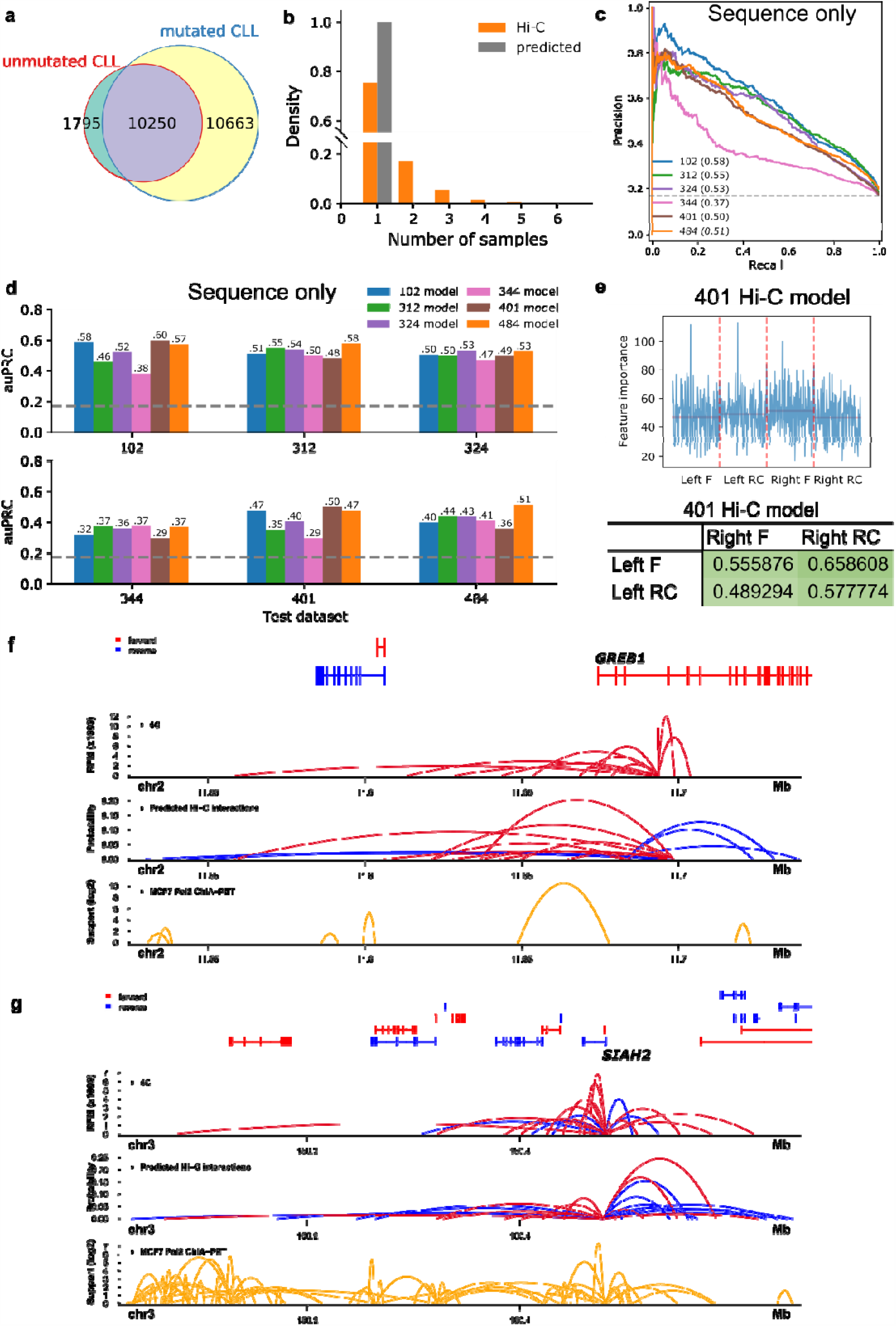
Performances of the sequence-based models in new CLL samples. **a**, Venn diagram of chromatin interactions identified by Juicer in unmutated and mutated CLL samples. **b**, Uniqueness analysis of real Hi-C and predicted Hi-C chromatin interactions in new CLL samples. Hi-C, real Hi-C interactions; predicted, predicted chromatin interactions using CLL 401 model. **c**, Precision-recall curves of the sequence-based models on distance-matched HiC datasets using only sequence features. **d**, Across-sample performances as measured by area-under precision-recall curve (auPRC) of the models on distance-matched HiC datasets using only sequence features. **e**, The importance scores of sequence features extracted from both directions (F: forward; RC: reverse complement) of the two anchors (left and right) by models trained on CLL 401 sample. The orange horizontal lines indicate average importance scores of the features from the strand of the anchor. Pearson correlations between feature importance scores of the two anchors are given in table. **f**, Validations of predicted chromatin interactions by 4C-seq at *GREB1* gene region in MCF-7 cells. In the predicted Hi-C interaction panel, only those interactions connected to *GREB1* promoter were shown. **g**, Validations of predicted chromatin interactions by 4C-seq at *SIAH2* gene region in MCF-7 cells. In the predicted Hi-C interaction panel, only those interactions connected to *SIAH2* promoter were shown.

After that, we trained gradient boosted trees with the corresponding extended datasets of the CLL samples. We observed that similar correlation of the important sequence features on different strands of the two anchors (Figure 6e; Supplementary Figure 8d-8e), although the within-sample and cross-sample auPRC were decreased (Supplementary Figure 8f-8g).

We also generated open chromatin pairs using ATAC-seq to train the gradient boosted trees (merging size: 3000 bp; extension size 1000 bp). Although the performances decreased compared with using Hi-C anchor region pairs as input, they were still higher than the random auPRC values (Supplementary Figure 8h-8k). We further used the 401 CLL sample model to predict chromatin interactions in MCF7 cells, as 401 CLL model have the highest within-sample and across-sample performance. The predicted interactions correlate quite well with the real 4C-seq interactions (Figure 6f-6g, Supplementary Figure 8l-8o, threshold = 0.016).

One question we asked was whether there is patient heterogeneity in Hi-C data. We first tried to associate the real and predicted Hi-C interactions with differentially expressed genes identified from RNA-seq data. The results showed that although the trend of different IFC scores could be observed, these differences were not significant (Supplementary Figure 8p-8q). We also observed that the Hi-C interactions and ATAC-seq peaks in the new CLL samples showed high patient heterogeneity (Supplementary Figure 8r). These patient heterogeneities may be a reason of the limited sample size in the IFC score analysis after we collapsed all six sample into mutated and unmutated categories (Supplementary Figure 8p-8q).

Taken together, our results demonstrate across-sample prediction capability for the ChINN model. In addition, we observed high patient heterogeneity in the new CLL samples, which may affect the predicted results and integrative analysis.

### Exploring chromatin interactions in a cohort of patient samples

Next, we used our machine learning method to predict chromatin interactions in a cohort of patient samples, and then analyzed the data. We applied the above models to 84 chronic lymphocytic leukemia (CLL) samples whose open chromatin profiles were available by ATAC-seq^24^.

A total of 48,443 CTCF-associated open chromatin interactions and 23,633 Pol2-associated open chromatin interactions were predicted based on the pooled open chromatin regions of all samples (Figure 7a). Pol2-associated chromatin interactions were better conserved across the CLL samples than CTCF-associated chromatin interactions (Figure 7b), which could be attributed to that open chromatin regions in the CLL samples that overlapped with GM12878 Pol2 peaks were better conserved than those overlapping with GM12878 CTCF peaks (Figure 7c). Using this set of ATAC-seq data in CLL samples, it was reported that regions with higher open chromatin signals in uCLL samples showed strong enrichment of binding sites of CTCF, RAD21 and SMC3^24^, which could also contribute to the high variability of CTCF chromatin interactions. Thus, we again observed extensive patient heterogeneity of CTCF and RNA Polymerase II-associated chromatin interactions in these clinical samples.

**Figure 7:**
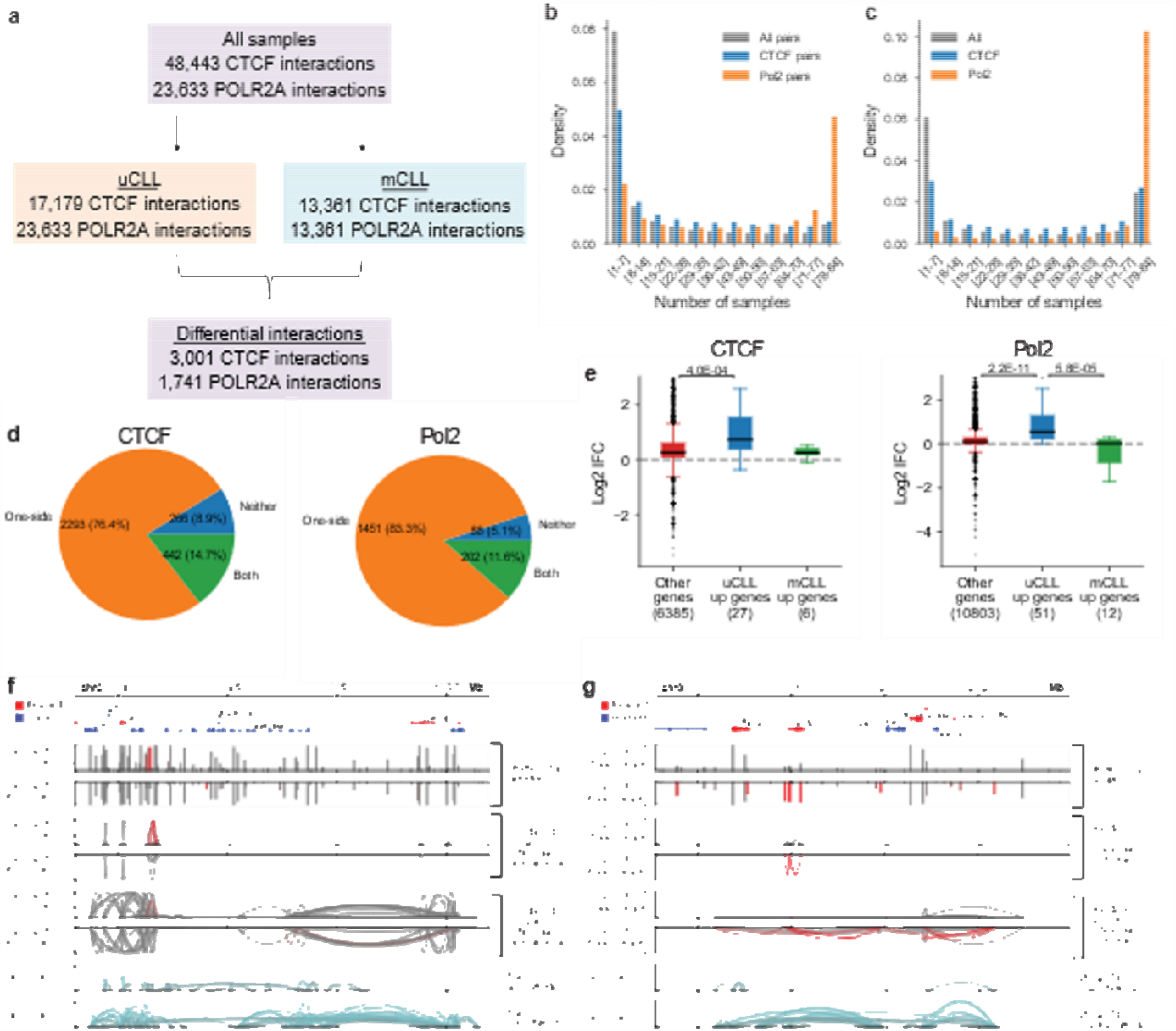
Predicted chromatin interactions in CLL samples. **a**, Summary of the predicted chromatin interactions in the 84 CLL samples and the differential chromatin interactions between uCLL and mCLL samples. **b**, Conservation analysis of predicted chromatin interactions in the CLL samples. All pairs: all possible pairs used for prediction. **c**, Uniqueness analysis of open chromatin regions that overlap with CTCF or Pol2 peaks from GM12878 cells in the CLL samples. All: all open chromatin regions. **d**, Distribution of differential CTCF and Pol2 chromatin interactions based on whether both anchors (Both), one anchor (One-side), or neither anchors (Neither) showed the same level of differences between uCLL and mCLL samples as the associated chromatin interaction. **e**, Association of differences in chromatin interactions between uCLL and mCLL samples with differentially expressed genes identified from a set of microarray samples. IFC: the fold change of the average number of chromatin interactions observed at the gene promoter in uCLL samples over that in mCLL samples. p-values were calculated using the Kruskal-Wallis test. **f-g**, Examples of genes, *ZBTB20* and *LPL*, whose different connectivity are associated with differences in distal regions. The red bars and curves indicate significantly different open chromatin regions and chromatin interactions based on Fisher’s Exact test.

When applying the GM12878 HiC model to the CLL samples, a total of 758,407 HiC-associated open chromatin interactions were predicted (Figure 8a). The phenomenon observed from the CTCF model also can be observed from the HiC model, for example, the chromatin interactions across the CLL samples and the overlapping peaks between CLL samples and GM12878 HiC peaks were not well conserved as that of Pol2 (Figure 8b-c). The predicted chromatin interactions by HiC model were also possible to separate mCLL and uCLL samples (Supplementary Figure 9a). Most differential chromatin interactions were associated with changes in the occurrence of one anchor (Figure 8d). Genes that were upregulated in uCLL were associated with uCLL-specific chromatin interactions (Figure 8e). In the set of differential chromatin interactions whose anchors did not have the same level of changes as the chromatin interactions themselves between the two subtypes, the rate of co-occurrences of the two anchors within the same sample and the levels in chromatin interactions could change (Supplementary Figure 9b). Examples of predicted chromatin interactions are shown in Figure 8f-g and Supplementary Figure 9e-h. Thus, we observed extensive patient heterogeneity of Hi-C predicted associated chromatin interactions in these clinical samples.

**Figure 8:**
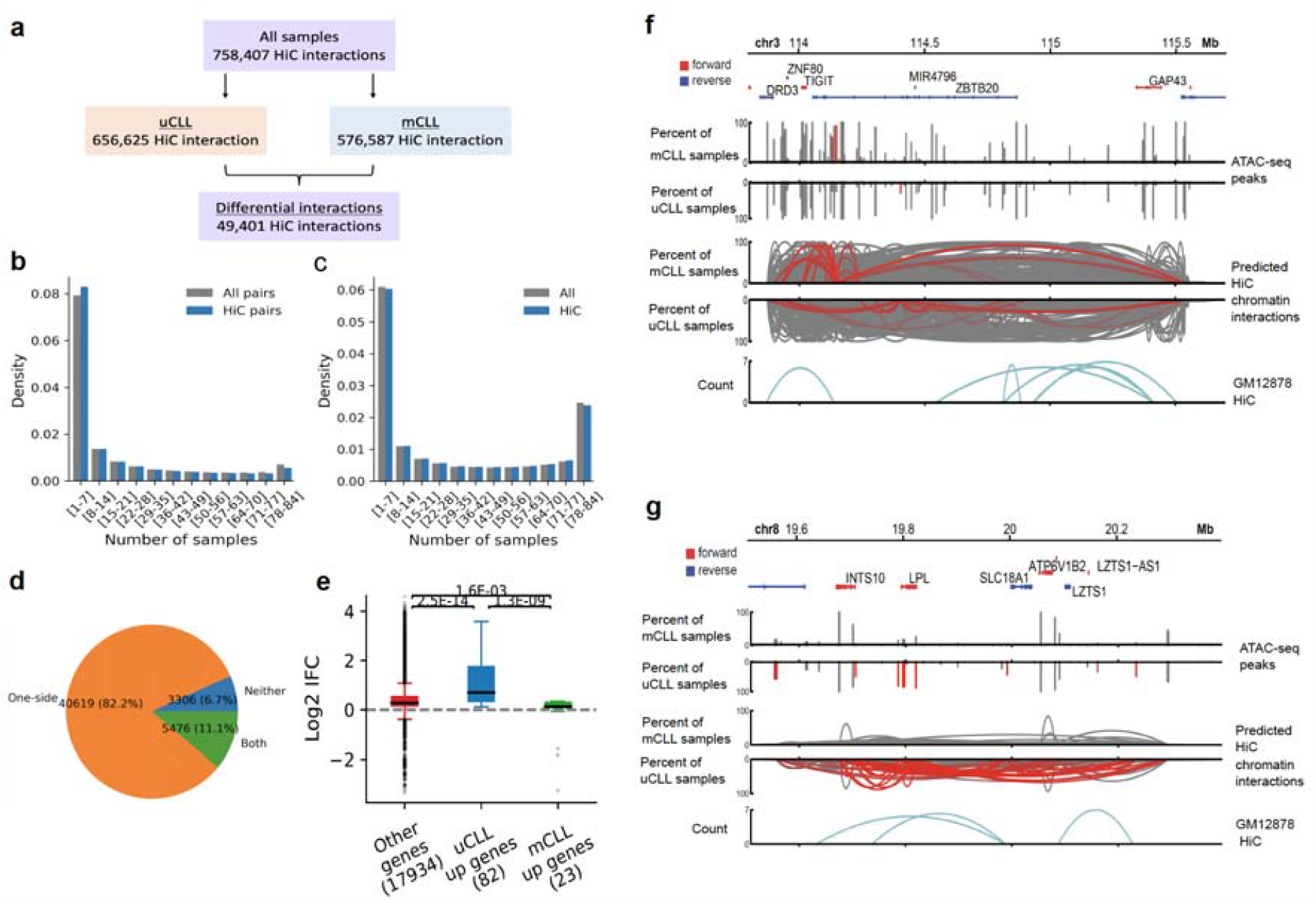
Predicted chromatin interactions in CLL samples using GM12878 HiC model. **a**, Summary of the predicted chromatin interactions in the 84 CLL samples and the differential chromatin interactions between uCLL and mCLL samples. **b**, Conservation analysis of predicted chromatin interactions in the CLL samples. All pairs: all possible pairs used for prediction. **c**, Uniqueness analysis of open chromatin regions that overlap with HiC peaks from GM12878 cells in the CLL samples. All: all open chromatin regions. **d**, Distribution of differential HiC chromatin interactions based on whether both anchors (Both), one anchor (One-side), or neither anchors (Neither) showed the same level of differences between uCLL and mCLL samples as the associated chromatin interaction. **e**, Association of differences in chromatin interactions between uCLL and mCLL samples with differentially expressed genes identified from a set of microarray samples. IFC: the fold change of the average number of chromatin interactions observed at the gene promoter in uCLL samples over that in mCLL samples. p-values were calculated using the Kruskal-Wallis test. **f-g**, Examples of genes, *ZBTB20* and *LPL*, whose different connectivity are associated with differences in distal regions. The red bars and curves indicate significantly different open chromatin regions and chromatin interactions based on Fisher’s Exact test.

Clinical samples differ from each other due to a wide variety of factors including different driver mutations and different underlying genetics and epigenetics of each patient. Here we asked whether the subtype of the CLL samples could be one factor giving rise to patient heterogeneity. The CLL samples could be divided into two subtypes based on IGHV mutation status: 34 IGHV-unmutated CLL (uCLL) samples and 50 IGHV-mutated CLL (mCLL) samples. IGHV mutation status is an important prognostic biomarker in CLL, with mCLL being less aggressive^24^.

Using the predicted chromatin interactions, it was possible to separate mCLL and uCLL samples (Supplementary Figure 10a). Variations in occurrences of chromatin interactions between the two subtypes of CLL were highly associated with variations in occurrences of anchor regions. Most differential ChIA-PET chromatin interactions were associated with changes in the occurrence of one anchor (Figure 7d). There was a small portion of differential chromatin interactions whose anchors did not have the same level of changes as the chromatin interactions themselves between the two subtypes. In this set of differential chromatin interactions, the rate of co-occurrences of the two anchors within the same sample could change, contributing to the levels of changes in chromatin interactions (Supplementary Figure 10b). With the GM12878 HiC model, we were also able to see differences in connectivity at transcription start sites associated with differences in the occurrences of the open chromatin regions at the transcription start sites (Supplementary Figure 9d).

Genes with higher expression in uCLL showed higher connectivity at the transcription start sites (Figure 7e, Supplementary Figure 10c, Figure 6e, Supplementary Figure 9c). The differences in connectivity at transcription start sites were associated with differences in the occurrences of the open chromatin regions at the transcription start sites between CLL subtypes (Supplementary Figure 10d), and also, differences in connectivity were sometimes associated with differences in distal interacting regions (Supplementary Figure 10e, Figure 7f). Examples of predicted chromatin interactions are shown at important CLL prognostic markers, such as *LPL* (Figure 7g), *ZAP70* (Supplementary Figure 10f), *ZNF667* (Supplementary Figure 10g), and *CD38* (Supplementary Figure 10h)^29-32^. Taken together, our results indicate that different subtypes show different profiles of chromatin interactions. Different subtypes may be a source of patient heterogeneity in clinical samples.

## Discussion

We described a convolutional neural network, ChINN, which can extract sequence features and be coupled to classifiers to predict chromatin interactions between open chromatin regions using DNA sequences and distance. This approach only requires the use of open chromatin data and showed good generalizability on the same type of chromatin interactions across different cell types. Thus, it has the potential to be applied to large sets of clinical samples with limited biological materials. In addition, CHINN can discover sequence features that are important for predicting chromatin interactions, including shared features such as the CTCF motif and cell-type specific features such as GATA3 binding motif in MCF-7, which is frequently mutated in breast cancer^33^.

We showed that at resolutions limited by the experimental techniques, chromatin interactions between open chromatin regions could be predicted from 1-dimensional functional genomics data with reasonable accuracy. In distance-controlled experiments, our prediction method using functional genomics data performed better on Pol2 chromatin interactions but worse on CTCF chromatin interactions compared to sequence-based CHINN. Such differences could be attributed to the lower functional genomic complexity at CTCF binding sites and functional genomic data might fail to capture the convergent CTCF binding motifs often observed at CTCF-mediated chromatin interactions.

On the other hand, Pol2 binding sites do not have such distinctive DNA motifs, making it harder to predict Pol2 binding sites^12, 13^ and consequently harder to predict Pol2-associated chromatin interactions from DNA sequences. However, Pol2 binding sites are usually occupied by many other transcription factors, making it easier to predict Pol2-associated chromatin interactions using functional genomic data.

The application of CHINN models with gradient boosted tree classifiers to a set of CLL ATAC-seq samples we were able to show that several of the predicted chromatin interactions could be validated by Hi-C. While there were also chromatin interactions that were predicted but not validated by Hi-C, our results showing that 4C could validate predicted chromatin interactions in MCF-7 cells that were not identified by Hi-C suggest that these so-called “false positives” might potentially be real chromatin interactions that were simply not captured by Hi-C due to limited sequencing depth of Hi-C libraries.

Additionally, application of ChINN models in CLL revealed that although there were chromatin interactions that were ubiquitous in all samples, there were a large number of patient-specific chromatin interactions and also chromatin interactions that were found in fewer than half the samples. One reason for these different chromatin interactions was due to different patient subtypes. We found systematic differences in chromatin interactions involving important CLL prognostic genes, such as *LPL* and *CD38*, between the IGHV-mutated and IGHV-unmutated subtypes. These results suggest that differences in chromatin interaction landscapes between CLL subtypes could have important functional implications in CLL biology.

Our observation of widespread patient heterogeneity in patient cancer samples highlights the need for precision medicine and the need to understand chromatin interactions in individual patient samples. Machine learning offers one way for us to predict chromatin interactions in a cost-effective manner. The CHINN method may be useful in the future in understanding chromatin interactions in large cohorts of clinical samples and identifying chromatin interaction-based biomarkers.

## Methods

We performed machine learning, Hi-C interaction analysis, ATAC-seq, RNA-seq and gene expression analyses as described in the following sections. A list of all libraries used and generated is provided in **Supplementary Table 6**.

### Machine learning of ChIA-PET data

The development of the sequence models was divided into three stages. In the first stage, the distance-matched datasets were used to train the models consist of convolutional neural network (feature extractor) with fully connected layers as the classier, as shown in Figure 2a. In the second and third stage, the feature extractors trained in the first stage were frozen and gradient tree boosting classifiers were used as classifiers. In the second stage, the gradient tree boosting classifiers were trained using the extended datasets. In the third stage, the gradient tree boosting classifiers were trained using all potential pairs of anchors generated from open chromatin data and annotated by existing ChIA-PET data. Thus, the final result was a program that took in a list of open chromatin regions and produced predictions of chromatin interactions between the open chromatin regions.

The feature extractors took DNA sequences of both anchors of a potential interacting pair as input. The classier then took the features generated by the feature extractor and optionally the distance between anchors as input and produced a probability score of interaction. More details can be found from Supplementary Methods1.

### Machine learning of Hi-C data from cell lines

We collected the Hi-C interactions from 8 cell lines, including GM12878, HeLaS3, HMEC, HUVEC, IMR90, K562, KBM7, and NHEK. The construction of machine learning model using Hi-C data from cell lines follows the same procedures as described in that of ChIA-PET data, where the positive data is annotated according to the Hi-C interactions.

### Machine learning of Hi-C data from clinical samples

We collected the Hi-C interactions from 6 CLL clinical samples, including CLL 102, CLL 312, CLL 324, CLL 344, CLL 401, and CLL 484. The construction of machine learning model using Hi-C data from cell lines follows the same procedures as described in that of ChIA-PET data, where the positive data is annotated according to the Hi-C interactions. The CLL 401 model was used in the across-sample prediction.

### Preparation of clinical samples

Chronic Lymphocytic Leukemia patient samples (either peripheral blood or bone marrow isolates) were obtained from the Leukemia Cell Bank at the National University Health System (NUHS) with patient consent, under Institute Review Board number H-20-022E. The CLL samples were either bone marrow aspirates (312,324,344,484, and 102) or peripheral blood (401). The samples were immediately frozen after collection and stored in liquid nitrogen until further use.

The samples were taken out of the liquid nitrogen and thawed by dipping in a beaker containing water at 37°C. Once the sample was thawed completely, the cells were immediately transferred to the 15ml falcon and resuspended in 10 ml PBS containing 2% fetal bovine serum (FBS) and 2mM EDTA. The cells were pelleted at 300 × g for 5 minutes at room temperature and resuspended in 5ml PBS containing 2% FBS and 2 mM EDTA. The cells were counted and checked for viability using Trypan Blue.

RNA and genomic DNA was isolated from the CLL patient samples using AllPrep DNA/RNA/miRNA universal kit (Qiagen) according to the to the manufacturer’s instructions. Briefly, cells lysate were homogenized by 21G needle and syringe together with lysis buffer and 1M DTT. After that, the homogenized lysate were transferred into AllPrep DNA mini spin column for genomic DNA extraction. The genomic DNA were then eluted by water and proceeded for the IGHV mutation test. The flow through after the AllPrep DNA mini spin column were then proceeded into RNease Mini spin column with on-column digestion for RNA extraction. The RNA were eluted in water and further sent for RNA-seq.

IGHV mutation test was performed following the method in Agathangelidis et al^34^. Briefly, IGHV-IGHD-IGHJ gene rearrangements were amplified by 5’ IGHV leader primers and 3’ IGHJ primers (primer sequences are provided in Supplementary Table 6) using genomic DNA (gDNA) from CLL patient samples. The PCR amplification was performed by PCR core kit (Qiagen). Final PCR products were imaged by agarose gel electrophoresis and purified by PCR purification kit (QIAGEN). Purified PCR products were confirmed through Sanger sequencing by 3’ IGHJ primers. The Sanger sequencing results were analysed by IMGT/V-QUEST tools35 to get the IGHV identity scores. If the identify score was larger than 98%, the CLL sample was considered unmutated sample while the score was lower than 98%, the CLL sample was considered as an mutated sample.

### In situ Hi-C

Hi-C libraries were prepared using the Arima Genomics kit (Arima Genomics, San Diego, CA) in conjunction with the Swift Biosciences Accel-NGS 2S Plus DNA Library Kit (Cat # 21024) and Swift Biosciences Indexing Kit (Cat # 26148) following the manufacturer’s recommendations. In brief, 1× 10^6^ cells were fixed with formaldehyde in the nucleus. Fixed cells were permeabilized using a lysis buffer and then digested with a restriction enzyme cocktail supplied in the Arima HiC kit. The resulting overhangs were filled in with biotinylated nucleotides followed by ligation. After ligation, crosslinks were reversed, and the DNA was purified from protein. Purified DNA was treated to remove biotin that was not internal to ligated fragments. Hi-C material was then sonicated using a Covaris Focused-Ultrasonicator M220 instrument to achieve 300-500 bp fragment sizes. The sonicated DNA was double-size selected using Ampure XP beads, and the sequencing libraries were generated using low input Swift Biosciences Accel-NGS 2S Plus DNA Library Kit (Cat # 21024) and Swift Biosciences Indexing Kit (Cat # 26148). The Hi-C libraries were loaded on an Illumina flow cell for paired-end 150-nucleotide read length sequencing on the Illumina HiSeq 4000 following the manufacturer’s protocols.

### Cell culture

MCF-7, a breast cancer cell line, was cultured in DMEM/F12 (Gibco) supplemented with 10% FBS and 1% penicillin-streptomycin and maintained at 37°C, 5% CO_2_ humidified incubator. Before 4C-seq assays, MCF-7 cells were grown in hormone-free media: they were washed with PBS twice to remove any residual FBS or growth factors and incubated in phenol red-free medium (Invitrogen/Gibco) supplemented with 10% charcoal-dextran stripped FBS (Hyclone) and 1% pencillin-streptomycin for a minimum of 72 hours. Hormone-depleted MCF-7 cells were then treated with oestrogen (Sigma) to a final concentration of 100 nM for 45 mins before 4C-seq assay. The control cells were treated with an equal volume and concentration of vehicle, ethanol (Sigma), for 45 min.

### Circular chromosome conformation capture (4C)

4C-seq assays were performed according to Splinter et al^68^ with slight modifications. Briefly, 4□×□10^7^ cells were cross-linked with 1% formaldehyde. The nuclei pellets were isolated by cell lysis with cold lysis buffer (10mM Tris-HCl, 10mM NaCl, 5mM EDTA, 0.5% NP 40) supplemented with protease inhibitors (Roche). First step digestion was performed overnight at 37°C with HindIII enzyme (NEB). Digestion efficiency was measured by RT-qPCR with HindIII site-specific primers. After confirmation of good digestion efficiency, DNA was ligated overnight at 16°C by T4 DNA ligase (Thermo Scientific) and de-crosslinked. Following de-crosslinking, DNA was extracted by phenol-chloroform and this is the 3C library. The DNA was then processed for second digestion with DpnII enzyme (NEB) overnight at 37°C. After final ligation, 4C template DNA was obtained, and the concentration was determined using Qubit assays (Thermo Scientific). The 4C template DNA was then amplified using specific primers with Illumina Nextera adapters and sent for sequencing on the MiSeq system. All the 4C genome coordinates are listed in Supplementary Table 6.

### RNA-seq

Total RNA was extracted from the CLL samples using the All Prep DNA/RNA kit (Qiagen). The RNA was quantified using the Qubit BR RNA Assay kit. RNA-seq libraries (strand specific and ribo zero) were constructed using Illumina Total RNA Prep kit (Illumina, San Diego, CA, USA) and sequenced 150 bases paired-end on the Illumina HiSeq 4000 following the manufacturer’s instruction.

### ATAC-seq

ATAC-seq library was prepared as described previously^36^. Briefly, 50,000 cells were lysed for nuclei isolation using ATAC-Resuspention Buffer containing 0.1% NP40, 0.1% Tween-20, and 0.01% Digitonin. Transposition reaction was performed for 30 min at 37C using Nextera DNA library preparation kit (NEB). Transposed fragments were amplified by eight PCR cycles for library preparation. Primer dimers and long DNA fragments were removed by AMPure XP beads purification step. DNA concentration was measured by Qubit fluorometric assay and library quality was determined by Bioanalyzer. The library was sequenced in Nextseq 500 76bp paired-end configuration using Illumina platform.

### Data deposition

The data for the RNA-Seq, ATAC-Seq and Hi-C data has been deposited with GEO accession number GSE163896. The 4C data has been deposited with GEO accession number GSE135052, and is publicly available.

## Data availability

All relevant data supporting the key findings of this study are available within the article and its Supplementary Information files or from the corresponding author on reasonable request.

## Code availability

The codes are freely available at: https://github.com/caofan/chinn.

## Acknowledgements

We would like to thank all members of the Fullwood Lab for helpful comments. We would like to thank the NUHS Leukemia Cell Bank for providing Chronic Lymphocytic Leukemia samples. This research is supported by the National Research Foundation (NRF) Singapore through an NRF Fellowship awarded to M.J.F (NRF-NRFF2012-054) and NTU start-up funds awarded to M.J.F. This research is supported by the RNA Biology Center at the Cancer Science Institute of Singapore, NUS, as part of funding under the Singapore Ministry of Education Academic Research Fund Tier 3 awarded to Daniel Tenen as lead PI with M.J.F as co-investigator (MOE2014-T3-1-006). This research is supported by a National Research Foundation Competitive Research Programme grant awarded to V.T. as lead PI and M.J.F. as co-PI (NRF-CRP17-2017-02). This research is supported by the National Research Foundation Singapore and the Singapore Ministry of Education under its Research Centres of Excellence initiative.

## Author Contributions

F.C and M.J.F. conceived of the research. F.C, Y.Z., Y.C.C. and M.J.F. contributed to the study design. F.C performed machine learning with CTCF and PolII ChIA-PET data. Y.Z. and Y.C.C. performed machine learning with Hi-C data. S.A. prepared Hi-C libraries from the CLL patient samples. Y.Z. performed genotyping of CLL patient samples. Y.P.L. performed 4C experiments. S.A. and V.T. designed ATAC-Seq experiments. S.A. performed ATAC-Seq experiments. CKK advised on machine learning analyses. F.C, Y.Z., Y.C.C. and M.J.F. reviewed the data and wrote the manuscript. All authors reviewed and approved of the manuscript.

## Notes

### Competing Interest Statement

The authors have declared no competing interest.

