## Supplementary Information for "Chromatin Interaction Neural Network (ChINN): A machine learning-based method for predicting chromatin interactions from DNA sequences"

2.2 Supplementary Figure 2: Convolutional layer 1 kernels of the five models that resemble known transcription factor binding motifs . . 15
2.3 Supplementary Figure 3: Sequence models on extended datasets . . 15
2.4 Supplementary Figure 4: HiC sequence models on extended datasets 15

4 Supplementary Figure 20

5 Supplementary Table 35

### 1 Methods

#### 1.1 Generation of positive and negative chromatin interactions

The GM12878 CTCF, GM12878 POLR2A, and Hela-S3 CTCF ChIA-PET datasets were downloaded from supplementary materials of Tang et al[1]. Alignment files of K562 POLR2A and MCF-7 POLR2A ChIA-PET datasets were obtained from Li et al[2] and processed as described in Li et al to call chromatin interactions.

Chromatin interactions that overlap with ENCODE blacklisted regions[3] at either anchor were discarded. Anchors of remaining chromatin interactions in each dataset were merged by allowing 500 bp gap[1]. The chromatin interactions in each dataset were clustered again using the merged anchors. Clustered chromatin interactions were filtered against DNase-seq data of the same cell line by requiring both anchors to overlap with DNase I hypersensitivity sites. The remaining clustered chromatin interactions were used as positive samples for each dataset.

For each dataset, corresponding negative samples were generated from four sources:

Source 1. Pairs of clustered chromatin interaction anchors that were not directly or indirectly connected were used as negative samples. We build networks of merged anchors where the merged anchors were nodes and the chromatin interactions were edges. Two nodes were considered as directly connected if there existed an edge between them. Two nodes were considered as indirectly connected if there existed a path of nodes and edges between the two nodes in the network.

Source 2. Random pairs of peaks from ChIP-seq data of the transcription factor from which the ChIA-PET library was built. The ChIP-Seq data and ChIA-PET data are from the same cell line. Peak pairs from CTCF ChIP-Seq data were used for CTCF ChIA-PET datasets and peak pairs from Pol2 ChIP-Seq data were used for POL2 ChIA-PET datasets. The peaks pairs were generated as follows:

(a) Transcription factor peaks that overlap with ENCODE blacklisted regions were filtered.

(b) From the remaining peaks, peaks that overlap with DNase I hypersensitivity regions were extracted.

(c) For each remaining peak, sample a size from a normal distribution. If the size of the peak is smaller than the sampled size, extend the peak to match the sampled size. The extended peaks were then merged. The mean and variance of the normal distribution were adjusted empirically to make the distribution of the sizes of merged peaks match the distribution of anchor sizes of the positive samples.

(d) All pairs of merged peaks with distance ranging from 5 kb to 2 Mb and were not directly or indirectly connected were kept.

Source 3. Random pairs of peaks from DNase-I data of the cell line from which the ChIA-PET library was built. The peak pairs were generated as follows:

(a) DNase-I peaks that overlap with ENCODE blacklisted regions were filtered.

(b) For each remaining peak, sample a size from a normal distribution. If the size of the peak is smaller than the sampled size, extend the peak to match the sampled size. The extended peaks are then merged. The mean and variance of the normal distribution were adjusted empirically to make the distribution of the sizes of merged peaks match the distribution of anchor sizes of the positive samples.

(c) All pairs of merged peaks with distance ranging from 5 kb to 2 Mb and were not directly or indirectly connected were kept. In addition, this set of negative samples were filtered against the negative samples generated using transcription factor peaks to reduce redundancy among negative samples.

Source 4. Pairs of clustered chromatin interaction anchors that were indirectly connected.

We generated two types of negative datasets: distance-matched negative datasets and extended negative datasets.

For each positive dataset, a distance-matched negative dataset was generated by sampling negative samples from the first three sources such that the positive-to-negative ratio was about 1:5 for every chromosome and the distance distributions were matched. We split the 5kb-2Mb distance range into 50 bins on the logarithm scale and counted the number of positive samples in each bin for each chromosome. Negative samples were then sampled such that there were about 5 negative samples for 1 positive sample in each bin of every chromosome. If the number of negative samples was less than 5 times of positive samples in a bin of a chromosome, more negative samples might be sampled from the same bin of another chromosome to make up the difference. Different types of negative samples were prioritized differently. Anchor pairs (Source 1) and transcription factor peak pairs (Source 2) were sampled first, and if there were not enough negative samples in these two sources, additional negative samples were sampled from DNase-I peak pairs (Source 3).

For each positive dataset, an extended negative dataset was generated by including pairs from Source 4, the corresponding distance-matched negative dataset, and the remaining negative samples from Source 1 and Source 2.

Samples on chromosomes 5 and 14 were used for validation. Samples on chromosomes 4, 7, 8, and 11 were used for test. The rest samples were used for training. The validation sets were used to guide the selection of the trained models. The test sets were used to obtained the final performance metrics and were strictly not used in training or selection of models.

Hi-C datasets were generated similarly as ChIA-PET. The cell line Hi-C interactions of GM12878, HeLa, HMEC, HUVEC, IMR90, K562, KBM7 and NHEK were downloaded from Rao et al[4].

#### 1.2 Overview of the functional genomic models

The functional genomic models were based on gradient tree boosting classifiers with features generated from functional genomic data such as ChIP-seq data and DNase-seq data. We and others have shown that many of the "state-of-art" methods based on functional genomic data had highly exaggerated performances and thus whether chromatin interaction could be accurately predicted from functional genomic data became unclear. This made our premise, chromatin interactions could be predicted from functional genomic data and functional genomic data could be predicted from DNA sequences, for developing predictive models based on DNA sequences broken. The purpose of developing the functional genomic models here was to confirm that chromatin interactions could be predicted from functional genomic data. For this purpose, we only trained and evaluated the models on distance-matched datasets.

#### 1.3 Generating features for functional genomic models

ChIP-seq peaks of histone marks and transcription factors were downloaded from ENCODE[3]. Histone marks and transcription factors that were common to K562, GM12878, and HelaS3 cells and DNase-seq peaks were used (Supplementary Table 1). For each sample, the feature values were calculated for both anchor independently. For an anchor $a$, let $F_a$ be its feature vector. Then the anchor's feature value of a transcription factor/histone modification/DNase $f$ is calculated as

$$F_{af} = \sum_i o_i^{af} v_i^f / s^a,$$

where $s^a$ is the size of the anchor, $o_i^{af}$ is the overlap size between the anchor $a$ and peak $i$ of factor $f$, and $v_i^f$ is the average signal value of peak $i$ of factor $f$. In short, a feature value of an anchor is the weighted sum of the signals of overlapping peaks of a factor, where the weights are the proportions of anchors in the overlaps.

The distance feature was calculated as the distance between the centers of two anchors.

We also evaluated models based on peak counts instead of signal values. When peak counts were used as features instead of signal values, we counted the peaks of a factor that overlap with an anchor and used the counts as feature values. In this case, a peak was only considered as overlapping with the anchor if the overlap size was at least 100 bp or at least 50% of the size of either the peak or the anchor.

#### 1.4 Training and evaluation of functional genomic models

We built functional genomic models for GM12878 CTCF, GM12878 POLR2A, HelaS3 CTCF, and K562 POLR2A datasets individually. The functional genomic models were constructed using gradient boosted trees from the XGBoost library[7]. Parameters used were: $max\_depth = 6$, $eta = 0.1$, $eval\_metric = logloss$. The models were trained for 1000 iterations with early stopping when the performances on validation sets were not improved for 40 consecutive iterations. The iteration of the models that produced best performance on validation sets were used as the final trained model for each dataset. The performance metrics were then obtained by applying the final trained models on the test sets.

#### 1.5 Overview of the sequence models

The development of the sequence models was divided into three stages. In the first stage, the distance-matched datasets were used to train the models consist of convolutional neural network (feature extractor) with fully-connected layers as the classifier, as shown in Figure 2a. In the second and third stage, the feature extractors trained in the first stage were frozen and gradient tree boosting classifiers were used as classifiers. In the second stage, the gradient tree boosting classifiers were trained using the extended datasets. In the third stage, the gradient tree boosting classifiers were trained using all potential pairs of anchors generated from open chromatin data and annotated by existing ChIA-PET data. Thus, the final result was a program that took in a list of open chromatin regions and produced predictions of chromatin interactions between the open chromatin regions.

The feature extractors took DNA sequences of both anchors of a potential interacting pair as input. The classifier then took the features generated by the feature extractor and optionally the distance between anchors as input and produced a probability score of interaction.

#### 1.6 Preparing input for sequence models

We used the human genome assembly *hg19* as our reference. The sequences of all anchors were extracted and converted to $4 \times L$ one-hot coded matrices, where $L$ was the length of the sequence. The four rows of the matrices represents the occurrences of nucleotides A, G, C, and T, respectively. Thus, nucleotide A was represented as $[1, 0, 0, 0]^T$, nucleotide G was represented as $[0, 1, 0, 0]^T$, nucleotide C was represented as $[0, 0, 1, 0]^T$, and nucleotide T was represented as $[0, 0, 0, 1]^T$. If N was encountered in the sequence, it was represented as $[0.25, 0.25, 0.25, 0.25]^T$. For each sequence, we split it into 1000 bp subregions with 500 bp overlap between consecutive subregions.

The distance feature was calculated as the distance between the centers of two anchors.

#### 1.7 The sequence model design considerations

A typical model based on convolutional neural network consists of two parts, the feature extracting convolutional layers (feature extractor) and the fully-connected layers (classifier) that aggregate the extract features and produce predictions or regressions. A convolutional layer is a stack of kernels, where each kernel is cross-correlated (convoluted) with the input independently, producing measures of the local similarities between the input and kernel. Non-linearities such as rectifier linear unit (ReLU) are applied on the outputs of convolutional layers to break the linearity. Convolutional layers are usually followed by max pooling layers to both reduce the spatial dimension of the output and allow small translations of the features on spatial dimensions. Max pooling layers take local maxima of convolutional outputs of each kernel independently. The output of the last convolutional layer is usually flattened and fed into the fully-connected layers to make predictions or regressions.

In this study, we used a weight-sum layer[8] after the last convolutional layer to obtain an aggregated score of each kernel in the last convolutional layer. The weighted-sum layer learns a weight distribution for each channel in the input independently and calculates the weighted sum of each channel along the spatial dimensions. Intuitively, the weighted-sum layer learns to weight the detected features at different spatial regions. For example, a weight-sum layer could have high weights for features detected near the center of the input sequence while penalizes features detected in surrounding regions by giving them negative weights. The weight-sum layer is followed by a non-linear activation function such as *tanh* or *sigmoid*. We used *tanh* for CTCF datasets and *sigmoid* for POLR2A datasets.

The sizes of chromatin interactions anchors vary substantially, ranging from kilo basepairs to tens of kilo basepairs. Thus, the network needs to accommodate the different anchor sizes. For each anchor, we split it into 1000bp subregions with 500bp overlap between consecutive subregions. The sequences of the subregions were fed into the convolutional layers independently. Following the non-linear function after weight-sum layer, maxima of each kernel across all subregions of an anchor were taken as the feature value for the anchor. Consequently, each anchor, regardless of its size, is represented by the same number of features.

#### 1.8 Mathematical formation of the components used

An anchor can be represented by a matrix $X$ of shape $N \times L \times C$, where $N$ is the number of subregions, $L$ is the length of the subregions and $C$ is the number of channels (or nucleotides in this application). For each kernel $k$ in a convolutional layer, let $S$ be the width of the kernel and $W^k$ of size $S \times C$ be the weight matrix of the kernel, $b^k$ be the bias, then

$$conv(X)_{n,m,k} = leakyReLU(\sum_{i=1}^{S}\sum_{j=1}^{C} X_{n,m+i,j}W^k_{i,j} + b^k);$$

$$leakyReLU(x) = \begin{cases} x & \text{if } x \geq 0 \\ \alpha x & \text{otherwise} \end{cases},$$

where $\alpha$ is a small constant and is set to be 1/5.5[9].

For a pooling layer, let $S$ be the size of the pool and $T$ be the stride, then for input $X$ of shape $N \times L \times C$,

$$pool(X)_{n,m,k} = \max_{i=1}^{S} X_{n,m*T+i,k}.$$

For input $X$ of $N \times L \times C$, the output of *weightsum* is of shape $N \times C$. For a weight-sum layer, let $W^{ws}$ be the weight matrix of shape $L \times C$, then

$$weightsum(X)_{n,k} = \sum_{i=1}^{L} X_{n,i,k}W^{ws}_{i,k}.$$

For POLR2A datasets, each value in the output of weight-sum layer is transformed by

$$sigmoid(x) = \frac{e^x}{1+e^x}.$$

For CTCF datasets, each value in the output of weight-sum layer is transformed by

$$tanh(x) = \frac{e^x - e^{-x}}{e^x + e^{-x}}$$

The features values of the anchor is then generated by taking the maximum of each kernel $k$ across all subregions by

$$maxfeature(X)_k = \max_{n=1}^{N} X_{n,k}.$$

#### 1.9 Training and evaluation of the sequence models on distance-controlled datasets

For each dataset, a model was trained using samples from its training set. Two rounds of training were performed for each model and each round was run for 50 epochs and early stopped if the loss on validation data was not improved in consecutive 10 epochs. The first round of training used larger learning rates and weighted positive samples by the negative-to-positive ratio in the training set, which is around 5. The second round of training used smaller learning rates. For CTCF models, the weights for positive samples were removed in the second round of training. The performances of the models on validation datasets were calculated after each epoch and the model at the epoch that produced the best performance were selected. The final performance metrics were then calculated using the test sets.

#### 1.10 Examining sequence features

The sequence patterns that kernels on the first and third convolutional layers detect were examined. For each kernel on first convolutional layer, as in DeepBind[10], we fed all test samples to the network and extracted a subsequence of each sample that pass the activation threshold of 0. If an anchor has multiple subsequences passing the activation threshold, the subsequence that has the highest activation is retained. Sequences that have no subsequences passing the activation threshold were discarded. The obtained subsequences for each kernel were then aligned to obtain a position-weight matrix for the kernel. The position-weight matrices of the kernels were compared against the transcription factor motif databases (HOCOMOCO version 11[11] and JASPAR core 2014 vertebrates collection[12]) using TomTom[13] with E-value threshold of 0.05.

For each kernel on the third convolutional layer, we generated an input matrix that maximize the output from the kernel. An input matrix was initialized randomly and fed into the network. The activation from the kernel under examination was obtained and the negation of its output was used as the loss to calculate the gradients on the input. The input matrix was then updated with stochastic gradient descent. The values of each column in the input matrix, which represents a base pair, were first clamped to the range of $[0, 1]$ and then normalized so that the sum of the values was 1. For each kernel, 20,000 iterations were performed.

The resulting input matrices were then used as position-weight matrices. As these position-weight matrices are 163 bp and TomTom takes long time to compare them to transcription factor motif databases, we split them into 30 bp smaller matrices with 20 bp stride. These smaller matrices were compared to transcription factor motif databases (HOCOMOCO version 11[11] and JASPAR core 2014 vertebrates collection[12]) using TomTom[13] with E-value threshold of 0.01.

#### 1.11 Training gradient boosted trees using extended datasets

The samples in the extended datasets were converted to matrices as described above. The convolutional layers trained using the distance-matched datasets were frozen and were used to produce feature vectors for the samples in the extended datasets. The generated feature vectors were used to train gradient tree boosting classifiers together with the distances between anchors. The splitting of samples into training, validation, and test sets were the same as described above.

We used gradient boosted trees from XGBoost library[7]. The *max_depth* setting was set to be 10, and trained for 1000 iterations and early stopped of if the loss on validation dataset is not improved in 40 consecutive iterations. The reported performance metrics were based on the test sets.

The classifiers on the five datasets achieved within-sample auPRCs ranging from 0.4 to 0.6 (Supplementary Figure 3a). If distance was not used, the performances of the classifiers dropped significantly (Supplementary Figure 3b). However, using distance alone could not predict chromatin interactions (Supplementary Figure 3c). These results suggested that the interplay between distance and sequence features were important in predicting chromatin interactions in general. The across-sample performances of the models on extended datasets were comparable to the corresponding within-sample performances, demonstrating the generalizability of the predictive models to new cell types using only sequence features and distance (Supplementary Figure 3d).

#### 1.12 CTCF-only models

We obtained all occurrences of CTCF motif in the genome using FIMO[14]. For each strand of each anchor, sum of matching scores of strand-specific occurrences of the CTCF motif within the anchor was calculated. The values were then normalized by the size of the anchors. Thus, for each chromatin interaction, a feature vector of four values were generated, each representing a strand of an anchor. Using these features together with distance, we trained another set of gradient boosted trees on the extended CTCF datasets.

#### 1.13 Training gradient boosted trees using pairs of open chromatin regions in GM12878

Two parameters, merging distance and extension size, were considered in determining how to generate anchors from DNase I hypersensitivity sites. As shown in Figure 4a, merging distance determines the gap allowed between neighboring peaks to merge them and extension size determines how much to extend on both ends of the merged peaks. To decide the two parameters, we used different combinations of the two parameters to process the DNase I peaks to generate anchors. Anchor pairs were then generated from the list of anchors such that the distance between centers of the two anchors were in the range of 5 kb to 2 Mb. Anchor pairs overlapping with ChIA-PET chromatin interactions were labelled as positive and the rest were labelled as negative. We then evaluated the performances of the models trained on extended datasets achieved on these anchor pairs for all combinations of the two parameters. The combination of the two parameters that achieved best area under ROC curve was selected. The reason to use auROC is because merging using different merging distances could lead to different number of anchor pairs generated and potentially different positive-to-negative ratios. Positive-to-negative ratios could affect auPRC measures and F1 measures as their baselines are sensitive to positive-to-negative ratio.

The DNase-I hypersensitivity sites were processed using the selected merging distance and extension size to produce anchors. Pairs of anchors were generated and labelled as described above. The anchor pairs were passed through the feature extractor to extract sequence features and generate feature vectors of sequence features. The sequence feature vectors and distance between anchors were then used to train gradient boosted tree classifiers. The parameters were then same as used for the extended datasets. The splitting of samples into training, validation and test sets followed the same rule as described above.

The trained classifiers produced a probability for each anchor pair. We noted that using 0.5 as the cutoff to call positive pairs did not produce optimal F1 scores. Thus, based on the F1 scores on the test sets, we selected the cutoffs that produced the best F1 scores for each dataset. The selected probability cutoffs were 0.22 for CTCF and 0.2 for POLR2A chromatin interaction predictions, respectively.

#### 1.14 4C-seq library generation

MCF-7, a breast cancer cell line, was cultured in DMEM/F12 supplemented with 10% FBS and 1% penicillin-streptomycin and maintained at 37°C, 5% $CO_2$ humidified incubator.

MCF-7 cells were grown in hormone-free media: they were washed with PBS and incubated in phenol red-free medium (Invitrogen/Gibco) supplemented with

10% charcoal-dextran stripped FBS (Hyclone) and 1% pencillin-streptomycin for a minimum of 72 hours. Hormone-depleted MCF-7 cells were treated with oestrogen to a final concentration of 100 nM for 45 mins before 4C assay. The control cells were treated with an equal volume and concentration of vehicle, ethanol (ET, Merck), for 45 min. In Supplementary Figure 5b-df, only 4C interactions in control cells were shown. The interactions detected by 4C were similar between oestrogen-treated and control MCF-7 cells (Supplementary Figure 5e-g).

For both MCF-7 and K562 cell lines, 4C-seq assays were performed as previously described in Splinter et al.[15] with slight modifications. Briefly, $4 \times 10^7$ cells were cross-linked with 1% formaldehyde and the nuclei pellets were isolated after cell lysis with cold lysis buffer supplemented with protease inhibitors. First step digestion was performed overnight at 37°C with HindIII enzyme. Digestion efficiency was measured by RT-qPCR with HindIII site-specific primers. After phenol-chloroform extraction, DNA was ligated overnight at 16°C by T4 DNA ligase. Following de-crosslinking, DNA was processed for second digestion with DpnII enzyme overnight at 37°C. After final ligation, 4C template DNA concentration was determined using fluorescence assay (picogreen, Invitrogen) and proceeded for library preparation for MiSeq sequencing using specific primers with Illumina Nextera adapters.

#### 1.15 4C-seq data analysis

Primer sequences at 5' ends of reads were trimmed using Tagdust2[16]. Extracted reads were then mapped to reference genome (hg19) using Bowtie2 (2.2.6)[17]. Unaligned reads were collected. The first 50 base pairs of the unaligned reads were extracted and realigned to the genome using Bowtie2 (2.2.6). The uniquely aligned reads (MAPQ$\geq$ 30) were combined. R Package r3Cseq[18] was used to call significant interactions using non-overlapping window approach. 5kb windows were used.

#### 1.16 RT-qPCR experiments

Total RNA were isolated from the cells using RNeasy Mini Kit (Qiagen) with on-column DNase digestion (Qiagen). 1$\mu$g of total RNA was then reverse transcribed to cDNA using the SuperScript III first-strand synthesis system using oligodT (Invitrogen). The expression levels of various genes were analysed by real-time PCR. Quantitative real-time PCR (qPCR) was performed on Applied Biosystems QuantStudio 3 Real-Time PCR system using SYBR Green PCR Master Mix and appropriate primers. The transcript levels of genes were analysed by $2^{-\Delta\Delta Ct}$ method.

#### 1.17 Analysis of chronic lymphocytic leukemia samples

ATAC-seq peaks of chronic lymphocytic leukemia (CLL) samples were collected from Gene Expression Omnibus (GSE81274). Peaks from the CLL samples were pooled and merged with merging distance of 3000 bp. The merged peaks were then extended by 1000 bp at both sides. All potential pairs of merged peaks that were on the same chromosome and had separation (as measured by the distance between centers of the two peaks) in the range of 5 kb to 2 Mb were used. The pairs were then fed into the CTCF and POLR2A CHINN models with gradient boosted trees trained on genome-wide open chromatin pairs to obtain probabilities of interaction. The probability cutoffs were 0.22 for CTCF chromatin interactions and 0.2 for POLR2A chromatin interactions, which were selected for the gradient boosted trees trained on genome-wide open chromatin pairs as described above. For each CLL sample, its CTCF chromatin interactions and POLR2A chromatin interactions were the predicted chromatin interactions whose both anchors overlap with peaks in the ATAC-seq library of the sample.

Random forest classifiers were trained using predicted CTCF and POLR2A chromatin interactions, respectively, to classify the samples into IGHV-mutated (mCLL) and IGHV-unmutated (uCLL) samples. Each sample is represented by a binary feature vector of the predicted chromatin interactions, where each value indicates whether the predicted chromatin interaction is present in the sample. We used the scikit-learn[19] implementation of random forest classifier with the parameter *n_estimator* set to 500. The performances were obtained by using 4-fold cross-validation with samples from the same patient always in the same group. Top 1000 important features for each trained random forest classifiers were then used for hierarchical clustering. Label-shuffled performances were obtained by training random forest classifiers on label-shuffled samples with the rest remain the same.

Differential chromatin interactions were obtained by using Fisher's Exact test to evaluate whether a chromatin interaction was more associated with any of the two CLL subtypes. The p-value threshold used was 0.01. The same Fisher's Exact test was applied to anchor peaks of the predicted chromatin interactions to test whether they were also more associated with any subtype of CLL. Differential chromatin interactions with both anchor peaks having p-values smaller than 0.01 were categorized as 'Both', those with only one anchor peak having p-value smaller than 0.01 were categorized as 'One-side', and those having neither anchor peak with p-value smaller than 0.01 were categorized as 'Neither'.

Two sets of CLL gene expression data were used: 1) a microarray dataset containing 76 mCLL samples and 58 uCLL samples from Gene Expression Omnibus (GSE69034); 2) a RNA-seq dataset containing 5 mCLL and 5 uCLL samples as an expression matrix downloaded from Gene Expression Omnibus (GSE81274). The RNA-seq dataset was from the same patient cohort as the ATAC-seq data. For the microarray dataset, limma[20] was used for differential gene expression analysis and genes with adjusted p-value smaller than 0.01 were selected. For the RNA-seq dataset, DESeq2[21] was used for differential gene expression analysis and genes with adjusted p-value smaller than 0.05 were selected.

Each predicted chromatin interaction $i$ and each open chromatin region $o$ was assigned a number $p_i^s$ or $p_o^s$ that indicates the percentage of samples of a subtype $s$ it appeared in. The connectivity $c_t^s$ of a transcription start site $t$ in a subtype $s$ was measured by taking the average of the percentages of samples in which its associated chromatin interactions appeared, which is

$$c_t^s = \sum_{i \in I} p_i^s / |I|$$

, where $I$ is the set of predicted chromatin interactions that overlapped with the transcription start site $s$. The presence of a transcription start site in a subtype was measured by the open chromatin region that was closest to the transcription start site. We then checked the fold changes of connectivity and presence of each transcription start site between uCLL and mCLL subtypes.

The Hi-C, RNA-seq, and ATAC-seq data of the six new CLL samples were deposited in GSE163896. For RNA-seq, kallisto (0.46.0)[22] was used to quantify the abundance of transcripts. The reference transcriptome used was Ensembl release 75. kallisto reports transcripts per million (TPM) for the transcripts. The TPM values were not nromalized across samples. For ATAC-seq, reads were aligned as paired-ends reads to hg19 genome assembly using BWA (0.7.17-r1188)[5]. The command used was bwa mem -M. Duplicates were marked and removed using MarkDuplicates function of Picard (2.22.2)[23] from GATK. Paired-end reads with either end unmapped were filtered using samtools (1.6)[24]. macs2 (2.2.6)[25] was used for peak calling with -BAMPE option. For Hi-C, BWA (0.7.17-r1188)[5] was used for aligning the paired-end reads to hg19 genome assembly. The command used was bwa mem -SP5M. The ENCODE HiC pipeline was used for filtering HiC alignments. Unmapped pairs and pairs with low quality mapping were removed. The pairs were deduplicated based on the mapping location of both ends. HiCCUPS from Juicer Tools (1.9.12)[6] was used for calling loops. Contact domains were detected using Arrowhead from Juicer Tools (1.9.12)[6].

### 2 Supplementary Figure Legends

#### 2.1 Supplementary Figure 1: Characteristics of positive and negative samples in distance-matched datasets

Supplementary Figure 1: Characteristics of positive and negative samples in distance-matched datasets. a, Comparison of the support of chromatin interactions based on whether both anchors (Both), only one anchor (XOR), or neither anchors (Neither) overlap withopen chromatin regions. The number in brackets indicates the number of chromatin interactions in the category. b, Distance distributions of positive and negative samples in distance-matched datasets.

#### 2.2 Supplementary Figure 2: Convolutional layer 1 kernels of the five models that resemble known transcription factor binding motifs

Supplementary Figure 2: Convolutional layer 1 kernels of the five models that resemble known transcription factor binding motifs

#### 2.3 Supplementary Figure 3: Sequence models on extended datasets

Supplementary Figure 3: Sequence models on extended datasets. a-c, Precision-recall curves of sequence models on extended datasets using (a) sequence features and distance; (b) only sequence features; and (c) only distance. d, Across-sample performancesof the sequence models on extended datasets. e-f, Top layer 3 kernels of CTCF models captured the CTCF motif. g, Performances of using only CTCF motif occurrences, motif orientations, and distance for prediction.

#### 2.4 Supplementary Figure 4: HiC sequence models on extended datasets

Supplementary Figure 4: HiC sequence models on extended datasets. a-c, Precision-recall curves of sequence models on extended datasets using (a) sequence features and distance; (b) only sequence features; and (c) only distance. d, Across-sample performances of the sequence models on extended datasets. e, Top layer 3 kernels of CTCF models captured the CTCF motif.

#### 2.5 Supplementary Figure 5: Selection of parameters for from-dnase models and validations by 4C-seq in MCF-7 cells

Supplementary Figure 5: Selection of parameters for from-dnase models and validations by 4C-seq in MCF-7 cells. a, auROC of models trained on extended datasets based on different combinations of merging distances and extension sizes. b, Validations of predicted chromatin interactions by 4C-seq at GREB1 gene gions in MCF-7 cells. c, Validations of predicted chromatin interactions by 4C-seq at SIAH2 gene gions in MCF-7 cells. d, Validations of predicted chromatin interactions by 4C-seq at EVOVL2 gene gions in MCF-7 cells. e-g: Comparison of 4C-seq chromatin interactions between oestrogen -treated and control MCF-7 cells at ELOVL2, GREB1 and SIAH2 gene.

#### 2.6 Supplementary Figure 6a-d: Selection of parameters for HiC models and validations by 4C-seq in MCF-7 cells

Supplementary Figure 6a-d: Selection of parameters for HiC models and validations by 4C-seq in MCF-7 cells. a, auROC of models trained on HiC extended datasets based on different combinations of merging distances and extension sizes. b, Validations of predicted chromatin interactions by 4C-seq at GREB1 gene gions in MCF-7 cells. c, Validations of predicted chromatin interactions by 4C-seq at SIAH2 gene gions in MCF-7 cells. d, Validations of predicted chromatin interactions by 4C-seq at EVOVL2 gene gions in MCF-7 cells.

#### 2.7 Supplementary Figure 7: The F-score with different thresholds for K562 and GM12878 HiC model on new CLL samples

Supplementary Figure 7: The F-score with different thresholds for K562 and GM12878 HiC model on new CLL samples. a, K562 HiC model. b, GM12878 HiC model.

#### 2.8 Supplementary Figure 8: Performances of the sequence-based and DNase models in new CLL samples.

Supplementary Figure 8: Performances of the sequence-based and DNase models in new CLL samples. a, Precision-recall curves of the sequence-based models on distance-matched Hi-C datasets using sequence features with distance. b, Across-sample performances as measured by area-under precision-recall curve (auPRC) of the models on distance-matched Hi-C datasets using sequence features with distance. c, Significant motifs identified from the CLL models by the convolutional layer 1 kernels. Identified motifs were ranked according to the p-value from the smallest (top) to the largest (bottom). E-value ¡ 0.01. d, The importance scores of sequence features extracted from both directions (F: forward; RC: reverse complement) of the two anchors (left and right) by models trained on CLL samples. The orange horizontal lines indicate average importance scores of the features from the strand of the anchor. e, Pearson correlations between feature importance scores of the two anchors of CLL samples. f, Performances of the sequence models on extended datasets compared with random model in CLL samples. g, Across-sample performances of the sequence models on extended datasets in CLL samples. h, Performances of the DNase model compared with random model in CLL samples. i, Confusion matrixes of the models in h, with threshold of 0.016. x-axis: ture label, y-axis: predicted label. 0: negative, 1: positive. j, Across-sample performances of the DNase model in CLL samples. k, Confusion matrixes of the results of using CLL 401 model to predict chromatin interactions in other samples with threshold of 0.016. x-axis: ture label, y-axis: predicted label. 0: negative, 1: positive. l-o, Validations of predicted chromatin interactions by 4C-seq at *ELOVL2*, *CAP2*, *P2RY2*, and *TP53* gene regions in MCF-7 cells. p-q, Association of differences in real Hi-C (p) and predicted Hi-C (q) chromatin interactions between uCLL and mCLL samples with differentially expressed genes from RNA-seq. IFC: the fold change of the average number of chromatin interactions observed at the gene promoter in uCLL samples over that in mCLL samples. r, ATAC-seq peaks, Hi-C interactions, and RNA-seq signal around KCNJ11 gene.

#### 2.9 Supplementary Figure 9: Analyses of predicted chromatin interactions in CLL samples using GM12878 HiC model

Supplementary Figure 9: Analyses of predicted chromatin interactions in CLL samples using GM12878 HiC model. a, Heatmap of hierarchical clustering of CLL samples based on the 1000 most important chromatin interactions of the HiC random forest classiers. Yellow indicates the presence of a chromatin interaction and black indicates the absence of a chromatin interaction. Red color represents mCLL samples and blue represents uCLL samples. b, Differences in agreement between anchors of differential HiC chromatin interactions in the 'Neither' category between mCLL and uCLL samples. c, Association of differences in chromatin interactions between uCLL and mCLL samples with differentially expressed genes identifed using RNA-seq data. IFC: the fold change of the average number of chromatin interactions observed at the gene promoter in uCLL samples over that in mCLL samples. p-values were calculated using Kruskal-Wallis test. d, The association between differences in chromatin interactions and differences in open chromatin peaks at gene promoter between uCLL and mCLL samples. Left: mutated, right: unmutated. PFC: the fold change of the proportions of samples the open chromatin peak at gene promoter is observed in uCLL samples over that in mCLL samples. The color barindicates the -log(pvalue) of the most significantly differential chromatin interaction at the promoter. e-h, Examples of genes whose different connectivity are associated with differences in distal regions. The red bars and curves indicate significantly different open chromatin regions and chromatin interactions based on Fisher's Exact test.

#### 2.10 Supplementary Figure 10: Analyses of predicted chromatin interactions in CLL samples

Supplementary Figure 10: Analyses of predicted chromatin interactions in CLL samples. a, Heatmaps of hierarchical clustering of CLL samples based on the 1000 most important chromatin interactions of the CTCF and Pol2 random forest classifiers, respectively. Yellow indicates the presence of a chromatin interaction and black indicates the absence of a chromatin interaction. Red color represents mCLL samples and blue represents uCLL samples. b, Differences in agreement between anchors of differential CTCF and Pol2 chromatin interactions in the 'Neither' category between mCLL and uCLL samples. c, Association of differences in chromatin interactions between uCLL and mCLL samples with differentially expressed genes identified using RNA-seq data. IFC: the fold change of the average number of chromatin interactions observed at the gene promoter in uCLL samples over that in mCLL samples. p-values were calculated using Kruskal-Wallis test. d, The association between differences in chromatin interactions and differences in open chromatin peaks at gene promoter between uCLL and mCLL samples. PFC: the fold change of the proportions of samples the open chromatin peak at gene promoter is observed in uCLL samples over that in mCLL samples. The color bar indicates the -log(pvalue) of the most significantly differential chromatin interaction at the promoter. e-h, Examples of genes whose different connectivity are associated with differences in distal regions. The red bars and curves indicate significantly different open chromatin regions and chromatin interactions based on Fisher's Exact test.

### 3 Supplementary Table Legends

#### 3.1 Supplementary Table 1: List of ChIP-seq peaks downloaded from ENCODE.

#### 3.2 Supplementary Table 2: Convolutional layer 1 kernels of the eight Hi-C models from cell lines that resemble known transcription factor binding motifs.

#### 3.3 Supplementary Table 3: The number of cell lines for each motif that was detected in eight Hi-C models

#### 3.4 Supplementary Table 4: Clinical characteristics of the CLL samples

#### 3.5 Supplementary Table 5: Hi-C libraries summary of CLL samples

#### 3.6 Supplementary Table 6: Primer information

### 4 Supplementary Figure

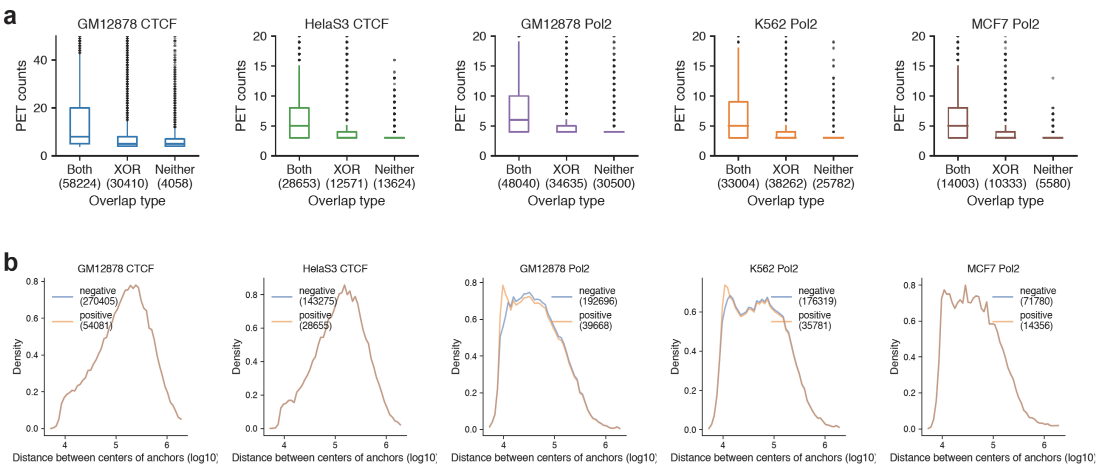

Supplementary Figure 1

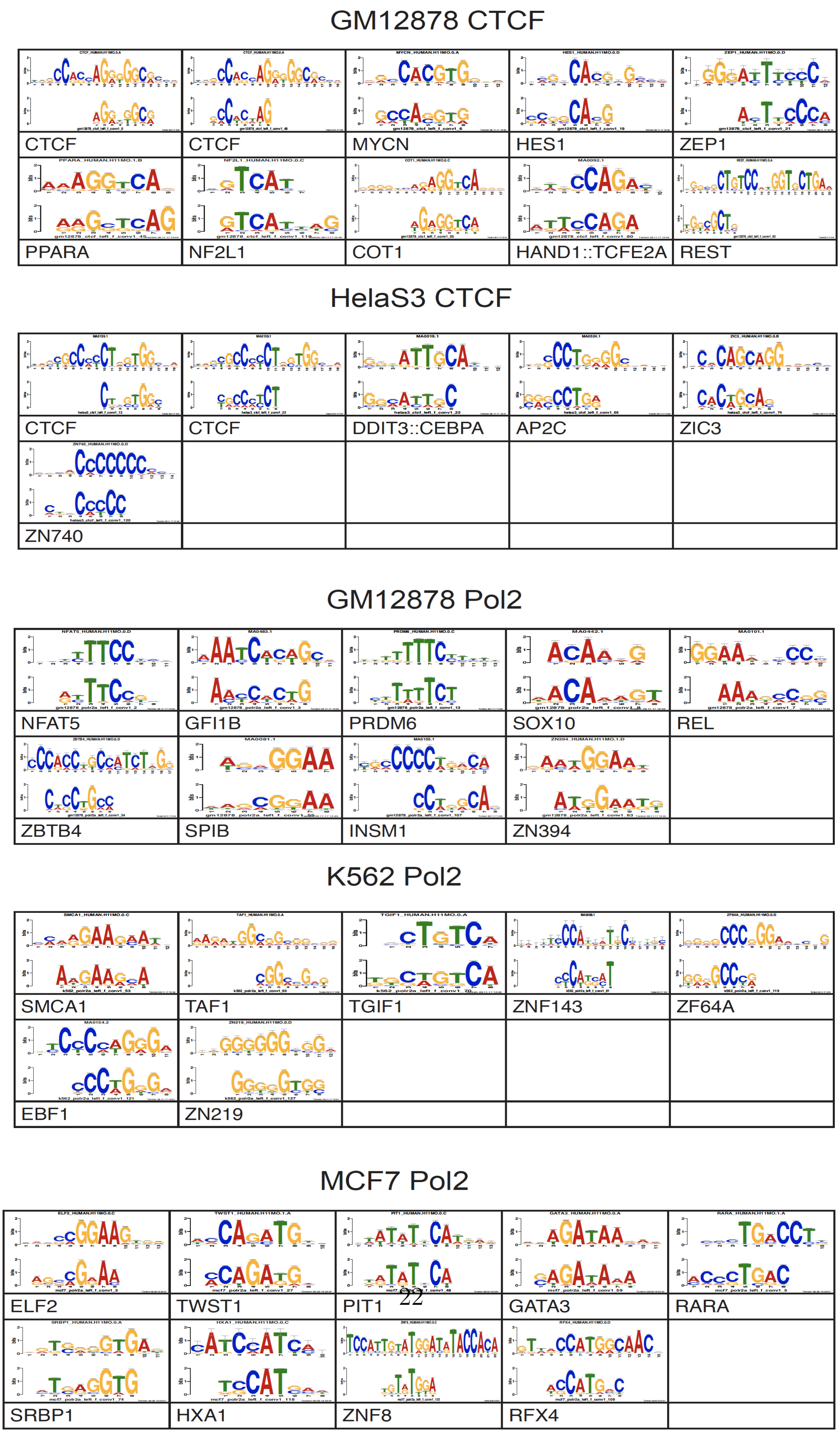

Supplementary Figure 2

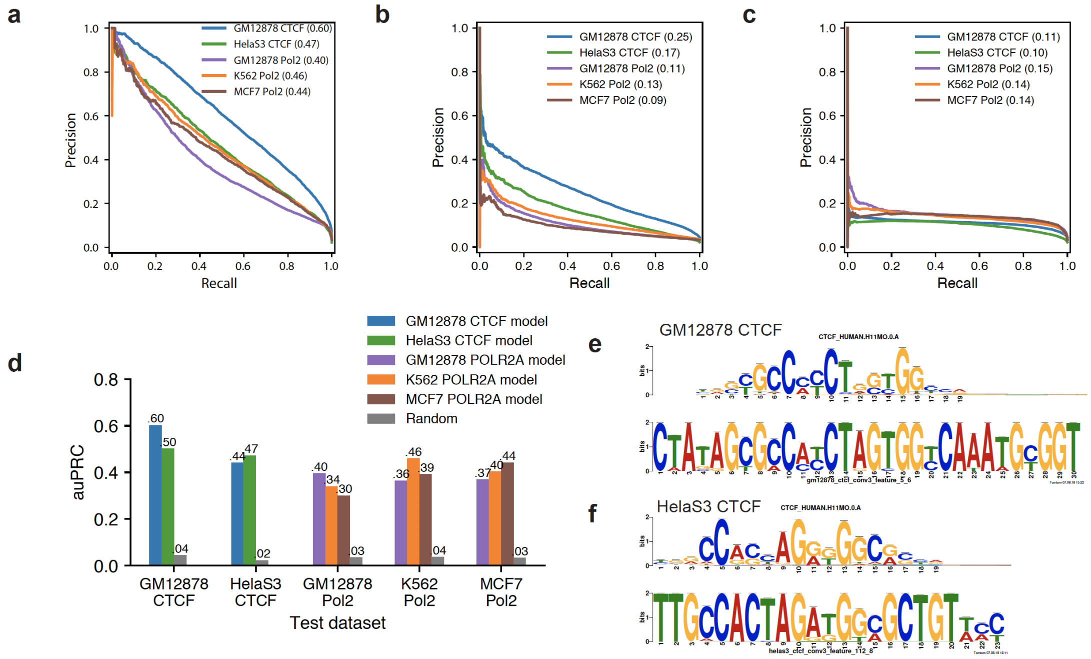

g

| auPRC | | Tested on | |
|---|---|---|---|
| | | GM12878 | HelaS3 |
| Trained on | GM12878 | 0.272 | 0.223 |
| | HelaS3 | 0.23 | 0.25 |

Supplementary Figure 3

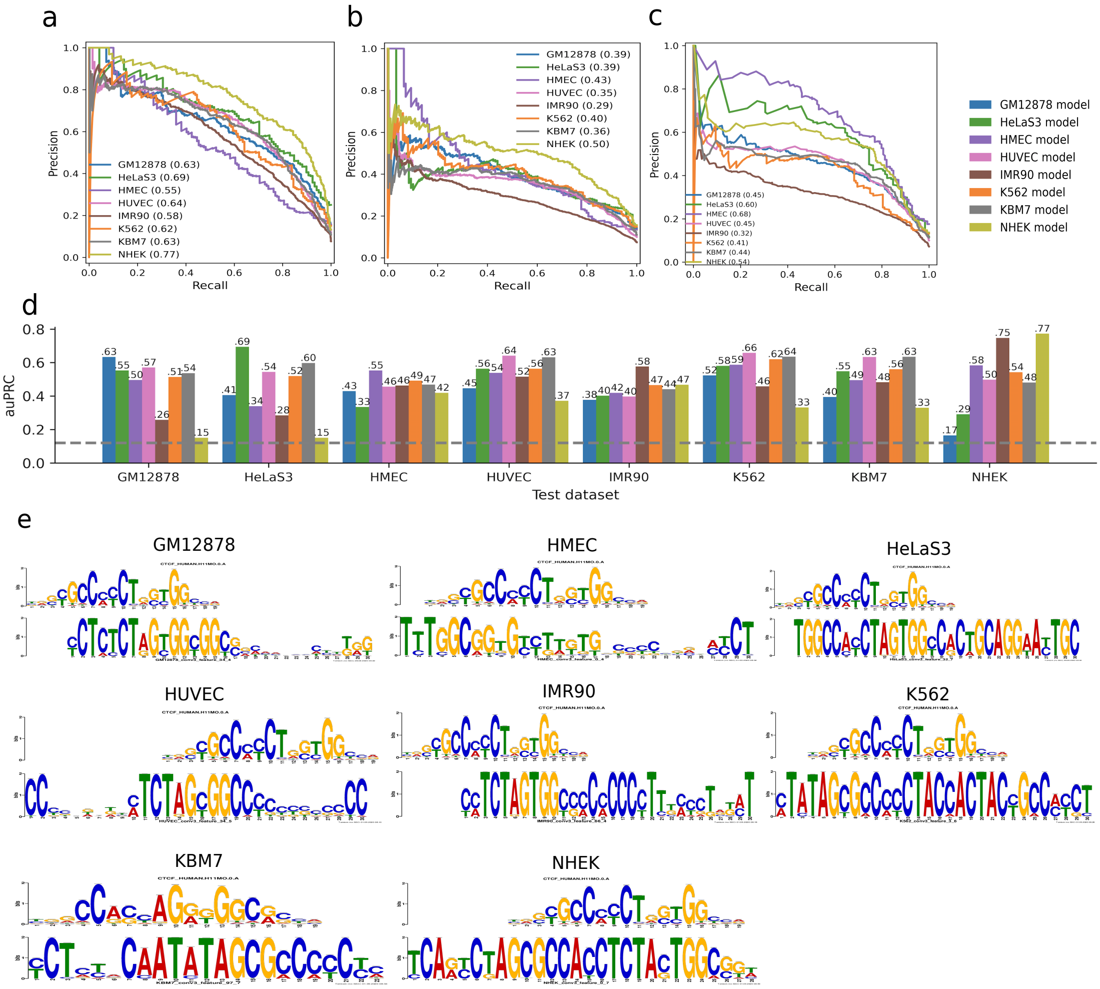

Supplementary Figure 4

**a**

| CTCF auROC | Extension size (bp) 500 | 1000 | 1500 | 2000 | 2500 | 3000 |
|---|---|---|---|---|---|---|
| Merging size (bp) 500 | 0.933 | 0.942 | 0.948 | 0.952 | 0.953 | 0.952 |
| 1000 | 0.944 | 0.946 | 0.95 | 0.954 | 0.954 | 0.954 |
| 1500 | 0.95 | 0.952 | 0.951 | 0.953 | 0.953 | 0.953 |
| 2000 | 0.953 | 0.955 | 0.954 | 0.954 | 0.953 | 0.953 |
| 2500 | 0.954 | 0.956 | 0.956 | 0.954 | 0.953 | 0.953 |
| 3000 | 0.955 | 0.957 | 0.956 | 0.955 | 0.954 | 0.952 |

| Pol2 auROC | Extension size (bp) 500 | 1000 | 1500 | 2000 | 2500 | 3000 |
|---|---|---|---|---|---|---|
| Merging size (bp) 500 | 0.926 | 0.932 | 0.934 | 0.934 | 0.934 | 0.934 |
| 1000 | 0.937 | 0.94 | 0.94 | 0.939 | 0.939 | 0.938 |
| 1500 | 0.943 | 0.946 | 0.946 | 0.945 | 0.945 | 0.944 |
| 2000 | 0.947 | 0.948 | 0.948 | 0.947 | 0.947 | 0.946 |
| 2500 | 0.953 | 0.954 | 0.953 | 0.953 | 0.952 | 0.951 |
| 3000 | 0.954 | 0.954 | 0.953 | 0.953 | 0.952 | 0.951 |

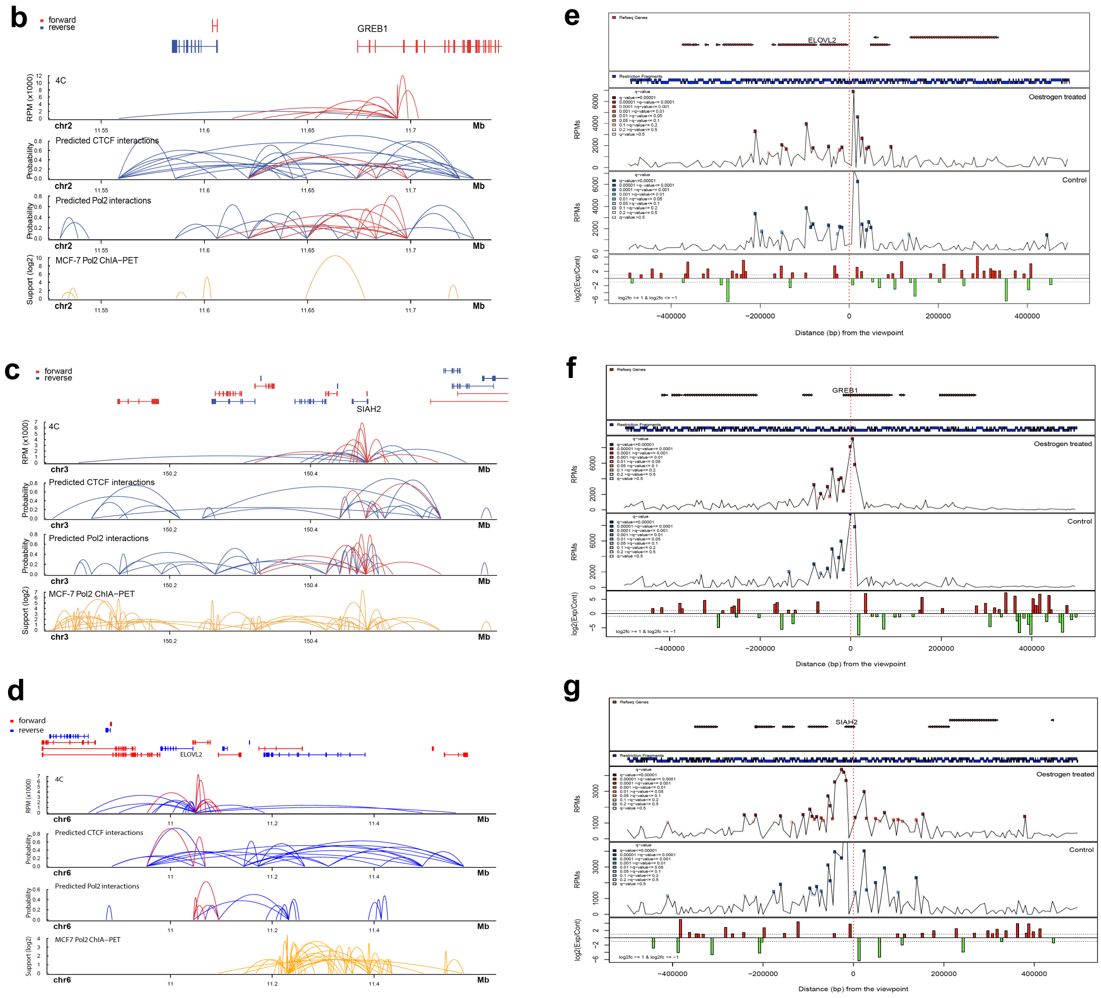

Supplementary Figure 5

**a**

| K562 auROC | | merge size 500 | 1000 | 2000 | 3000 | 4000 | 5000 |
|---|---|---|---|---|---|---|---|
| extension size | 500 | 0.8554 | 0.8549 | 0.7691 | 0.8582 | 0.8594 | 0.8543 |
| | 1000 | 0.8668 | 0.8654 | 0.8657 | 0.8693 | 0.8679 | 0.8652 |
| | 2000 | 0.8743 | 0.8769 | 0.8795 | 0.8765 | 0.8766 | 0.8747 |
| | 3000 | 0.8732 | 0.8809 | 0.8816 | 0.8775 | 0.8772 | 0.8758 |

| GM12878 auROC | | merge size 500 | 1000 | 2000 | 3000 | 4000 | 5000 |
|---|---|---|---|---|---|---|---|
| extension size | 500 | 0.8157 | 0.8129 | 0.8192 | 0.821 | 0.8257 | 0.8314 |
| | 1000 | 0.8061 | 0.8027 | 0.8069 | 0.8083 | 0.8159 | 0.8214 |
| | 2000 | 0.778 | 0.7778 | 0.7895 | 0.7904 | 0.7978 | 0.8039 |
| | 3000 | 0.7502 | 0.7477 | 0.7551 | 0.758 | 0.762 | 0.7687 |

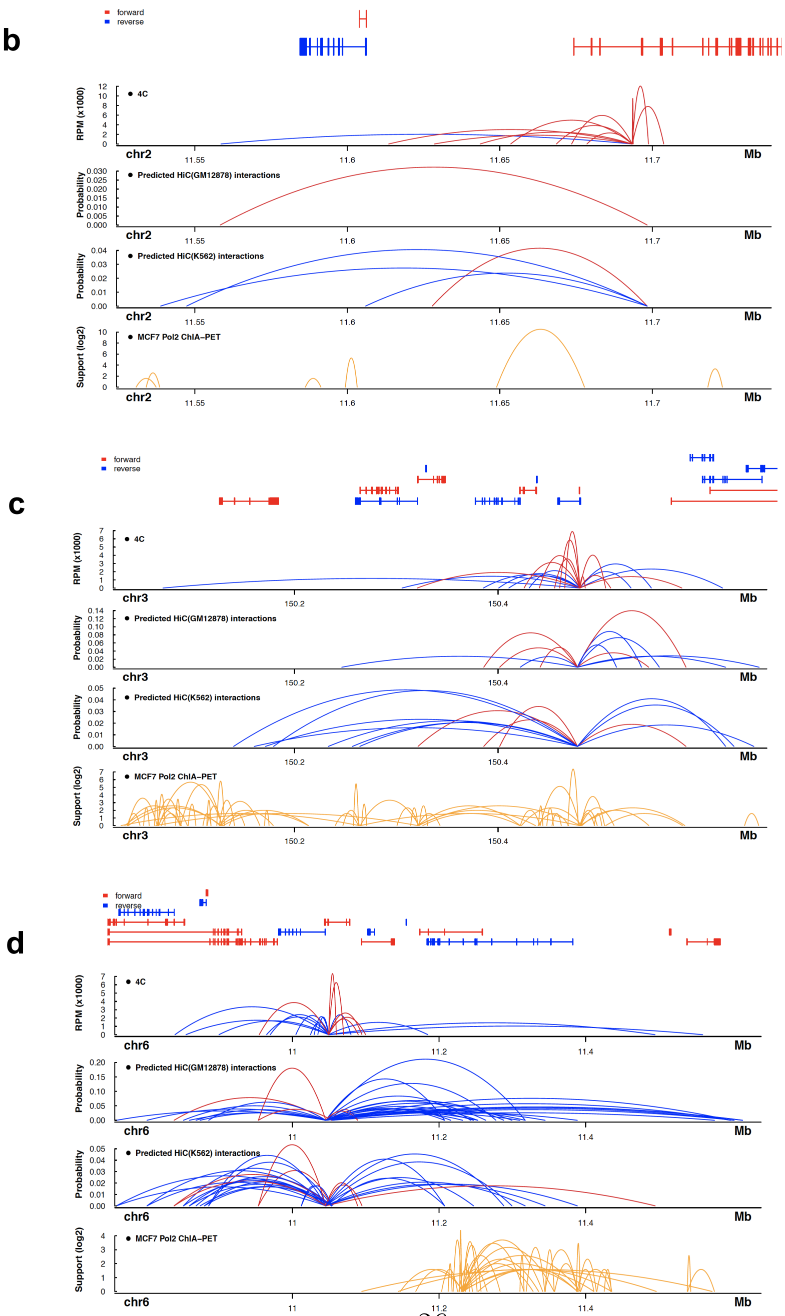

Supplementary Figure 6

**a**

| K562 F-score / cll sample | threshold 0.5 | 0.4 | 0.3 | 0.2 | 0.1 | 0.03 | 0.025 | 0.02 | 0.017 | 0.016 | 0.015 | 0.014 | 0.01 |
|---|---|---|---|---|---|---|---|---|---|---|---|---|---|
| 102 | 0.0017 | 0.0034 | 0.0076 | 0.0241 | 0.0929 | 0.2989 | 0.3169 | 0.3297 | 0.3351 | 0.3351 | 0.3325 | 0.33 | 0.3182 |
| 312 | 0.0005 | 0.0021 | 0.0077 | 0.036 | 0.1315 | 0.3618 | 0.3811 | 0.3964 | 0.4094 | 0.4122 | 0.4121 | 0.4101 | 0.3953 |
| 324 | 0.0013 | 0.0019 | 0.0076 | 0.0351 | 0.1409 | 0.3797 | 0.3979 | 0.419 | 0.4223 | 0.4229 | 0.422 | 0.4197 | 0.403 |
| 344 | 0.0006 | 0.0006 | 0.005 | 0.0186 | 0.0726 | 0.2426 | 0.2669 | 0.2947 | 0.3067 | 0.3138 | 0.3157 | 0.3162 | 0.322 |
| 401 | 0.0009 | 0.0015 | 0.0043 | 0.0177 | 0.0773 | 0.2516 | 0.2704 | 0.2845 | 0.2866 | 0.2894 | 0.29 | 0.2922 | 0.2894 |
| 484 | 0.0007 | 0.0014 | 0.0036 | 0.0146 | 0.0678 | 0.2567 | 0.2859 | 0.3132 | 0.3316 | 0.3394 | 0.3449 | 0.3484 | 0.3625 |

**b**

| GM12878 F-score / cll sample | threshold 0.5 | 0.4 | 0.3 | 0.2 | 0.1 | 0.03 | 0.028 | 0.027 | 0.025 | 0.024 | 0.02 | 0.015 | 0.01 |
|---|---|---|---|---|---|---|---|---|---|---|---|---|---|
| 102 | 0.0013 | 0.0025 | 0.0138 | 0.0565 | 0.2025 | 0.3843 | 0.385 | 0.3846 | 0.3804 | 0.3779 | 0.3659 | 0.3536 | 0.3338 |
| 312 | 0.0005 | 0.0036 | 0.0173 | 0.0705 | 0.2388 | 0.3841 | 0.3812 | 0.3816 | 0.382 | 0.3818 | 0.3696 | 0.3595 | 0.3406 |
| 324 | 0.0013 | 0.0044 | 0.0206 | 0.0813 | 0.2656 | 0.4455 | 0.4465 | 0.4466 | 0.4484 | 0.4473 | 0.4354 | 0.4089 | 0.3685 |
| 344 | 0 | 0.0023 | 0.0134 | 0.0478 | 0.163 | 0.2958 | 0.3027 | 0.3034 | 0.3063 | 0.3087 | 0.3143 | 0.3179 | 0.318 |
| 401 | 0.0006 | 0.0025 | 0.0152 | 0.0535 | 0.1674 | 0.335 | 0.3365 | 0.3373 | 0.3368 | 0.3354 | 0.3284 | 0.326 | 0.319 |
| 484 | 0.0007 | 0.0036 | 0.0126 | 0.0453 | 0.1546 | 0.3328 | 0.3417 | 0.3448 | 0.3518 | 0.355 | 0.37 | 0.3856 | 0.3931 |

Supplementary Figure 7

**a** **b**

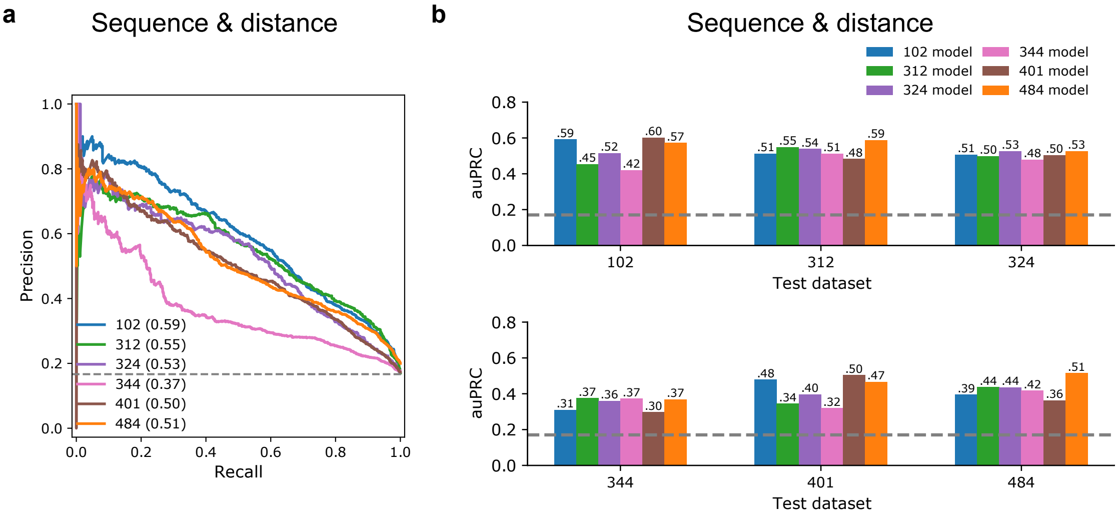

**c**

| | 102 | 312 | 324 | 344 | 401 | 484 |
|---|---|---|---|---|---|---|
| Motif | CTCF | CTCF | CTCF | CTCF | CTCF | CTCF |
| | CTCFL | CTCFL | CTCFL | CTCFL | CTCFL | CTCFL |
| | FOS | FOS | FOS | FOS | FOS | FOS |
| | JUNB | JUNB | JUNB | JUNB | JUNB | JUNB |
| | FOSL1 | FOSL1 | FOSL1 | FOSL1 | FOSB | FOSL1 |
| | JUN | JUN | JUN | JUN | FOSL1 | JUN |
| | JUND | JUND | JUND | JUND | JUN | JUND |
| | NFE2 | FOSL2 | FOSB | FOSL2 | JUND | FOSL2 |
| | FOSB | FOSB | FOSL2 | FOSB | NFE2 | FOSB |
| | BACH2 | NF2L2 | NFE2 | NFE2 | BACH2 | NFE2 |
| | MAFK | NFE2 | ZN436 | NF2L2 | MAFK | NF2L2 |
| | FOSL2 | BACH2 | NF2L2 | BACH2 | BACH1 | BACH2 |
| | BACH1 | MAFK | BACH2 | ZN436 | FOSL2 | BACH1 |
| | NF2L2 | BACH1 | MAFK | MAFK | ZN436 | MAFK |
| | ZN436 | ZN436 | BACH1 | BACH1 | NF2L2 | ZN436 |
| | INSM1 | TFE3 | BMAL1 | BMAL1 | INSM1 | IRF4 |
| | | TFEB | CR3L1 | CR3L1 | | IRF8 |
| | | USF1 | HES5 | HES5 | | CR3L1 |
| | | USF2 | HES7 | HES7 | | MYCN |
| | | CR3L1 | HEY1 | HEY1 | | BMAL1 |
| | | HES5 | HEY2 | MITF | | HES5 |
| | | HES7 | MITF | MYC | | HES7 |
| | | HEY1 | MYC | MYCN | | HEY1 |
| | | MITF | MYCN | TFE3 | | HEY2 |
| | | MYC | TFE3 | USF1 | | MITF |
| | | MYCN | USF1 | HEY2 | | MXI1 |
| | | BMAL1 | INSM1 | TFEB | | MYC |
| | | MAFG | MAX | USF2 | | TFE3 |
| | | INSM1 | MXI1 | MXI1 | | USF1 |
| | | | TFEB | MAX | | |
| | | | USF2 | | | |

Supplementary Figure 8 a-c

**d**

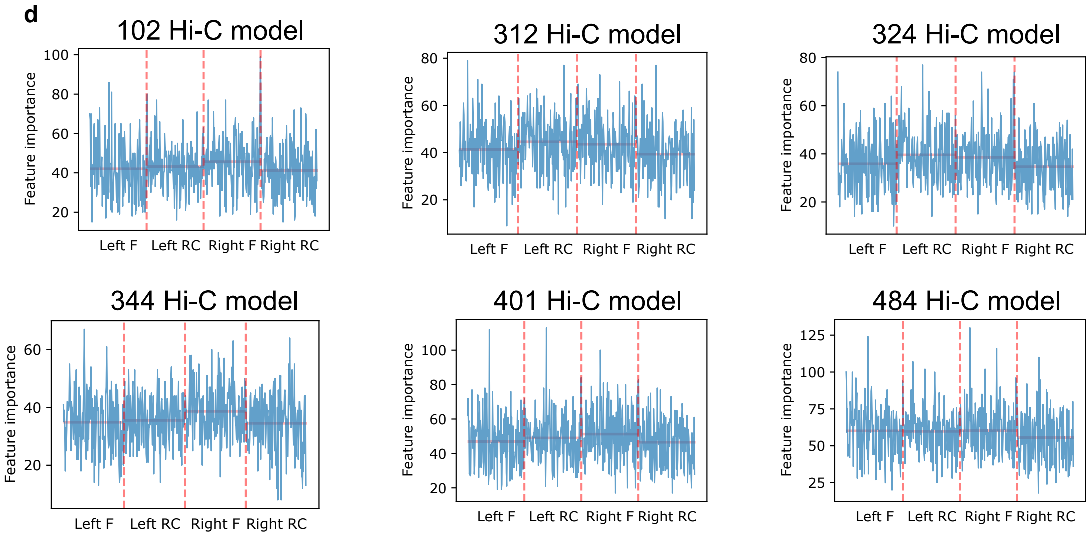

**e**

102 Hi-C model

| | Right F | Right RC |
|---|---|---|
| Left F | 0.562181 | 0.584517 |
| Left RC | 0.510532 | 0.480287 |

312 Hi-C model

| | Right F | Right RC |
|---|---|---|
| Left F | 0.380838 | 0.52689 |
| Left RC | 0.452727 | 0.366889 |

324 Hi-C model

| | Right F | Right RC |
|---|---|---|
| Left F | 0.444549 | 0.51084 |
| Left RC | 0.472598 | 0.387157 |

344 Hi-C model

| | Right F | Right RC |
|---|---|---|
| Left F | 0.384301 | 0.571981 |
| Left RC | 0.386465 | 0.416936 |

401 Hi-C model

| | Right F | Right RC |
|---|---|---|
| Left F | 0.555876 | 0.658608 |
| Left RC | 0.489294 | 0.577774 |

484 Hi-C model

| | Right F | Right RC |
|---|---|---|
| Left F | 0.606085 | 0.720893 |
| Left RC | 0.58604 | 0.628004 |

Supplementary Figure 8 d-e

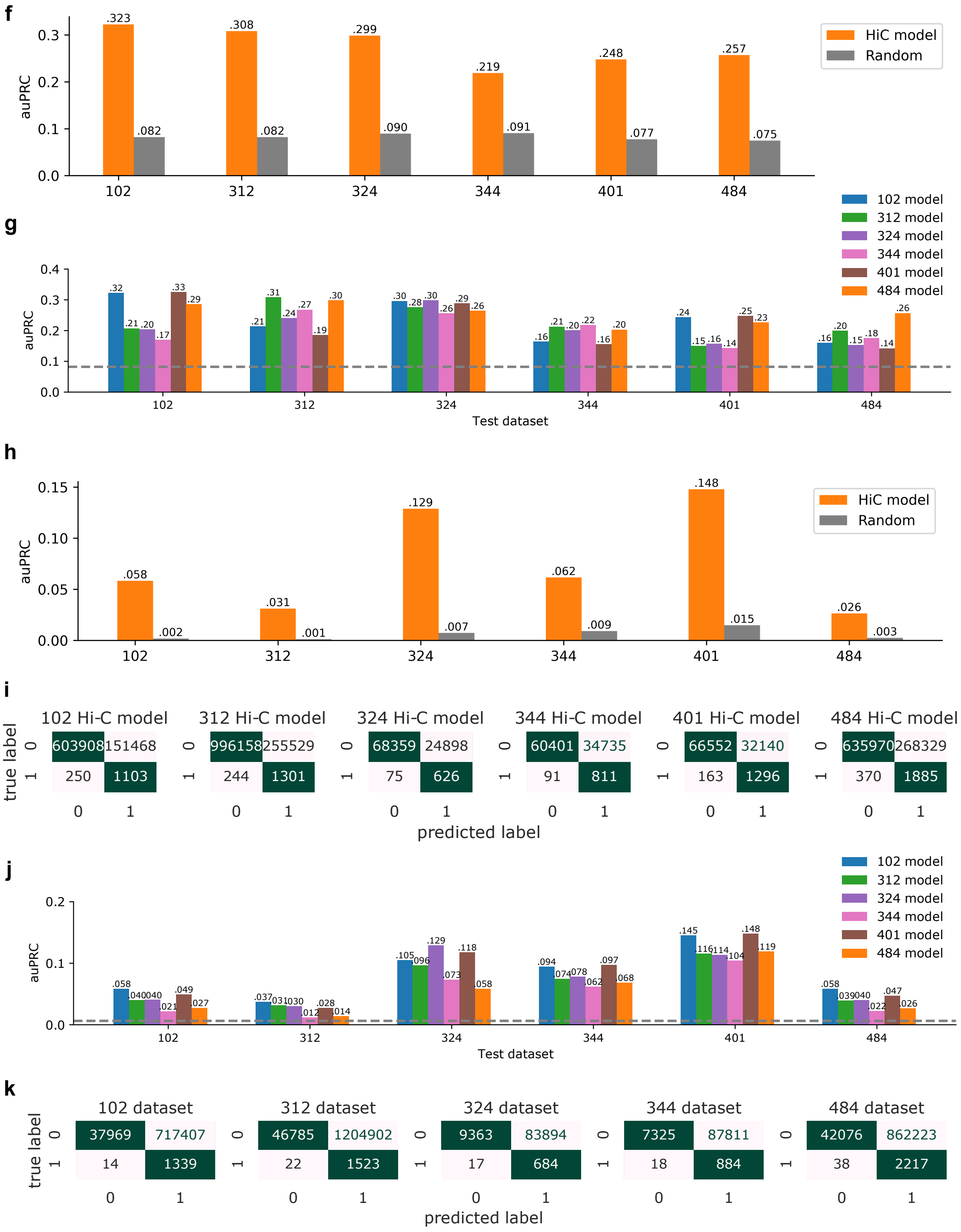

Supplementary Figure 8 f-k

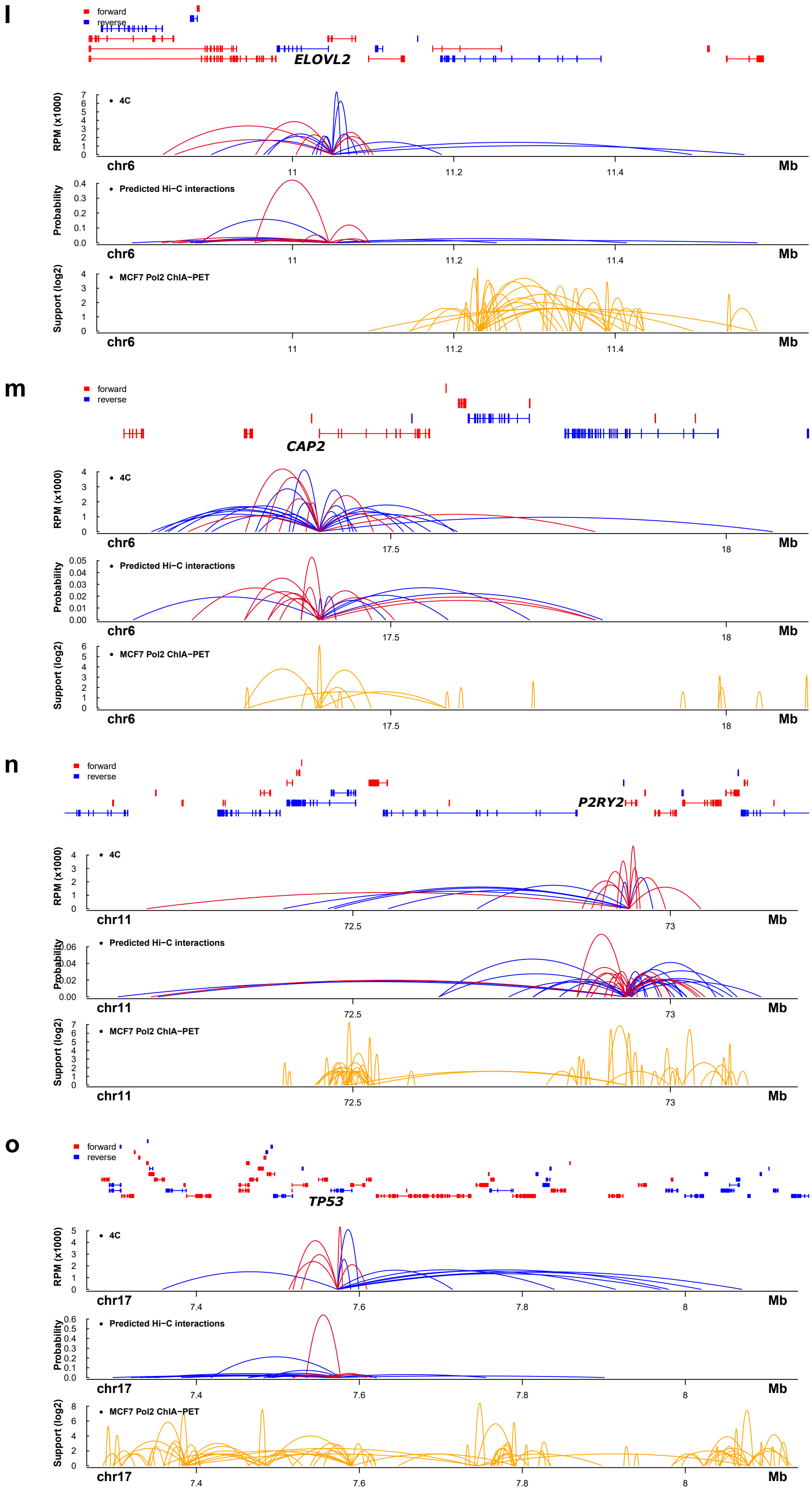

Supplementary Figure 8 l-o

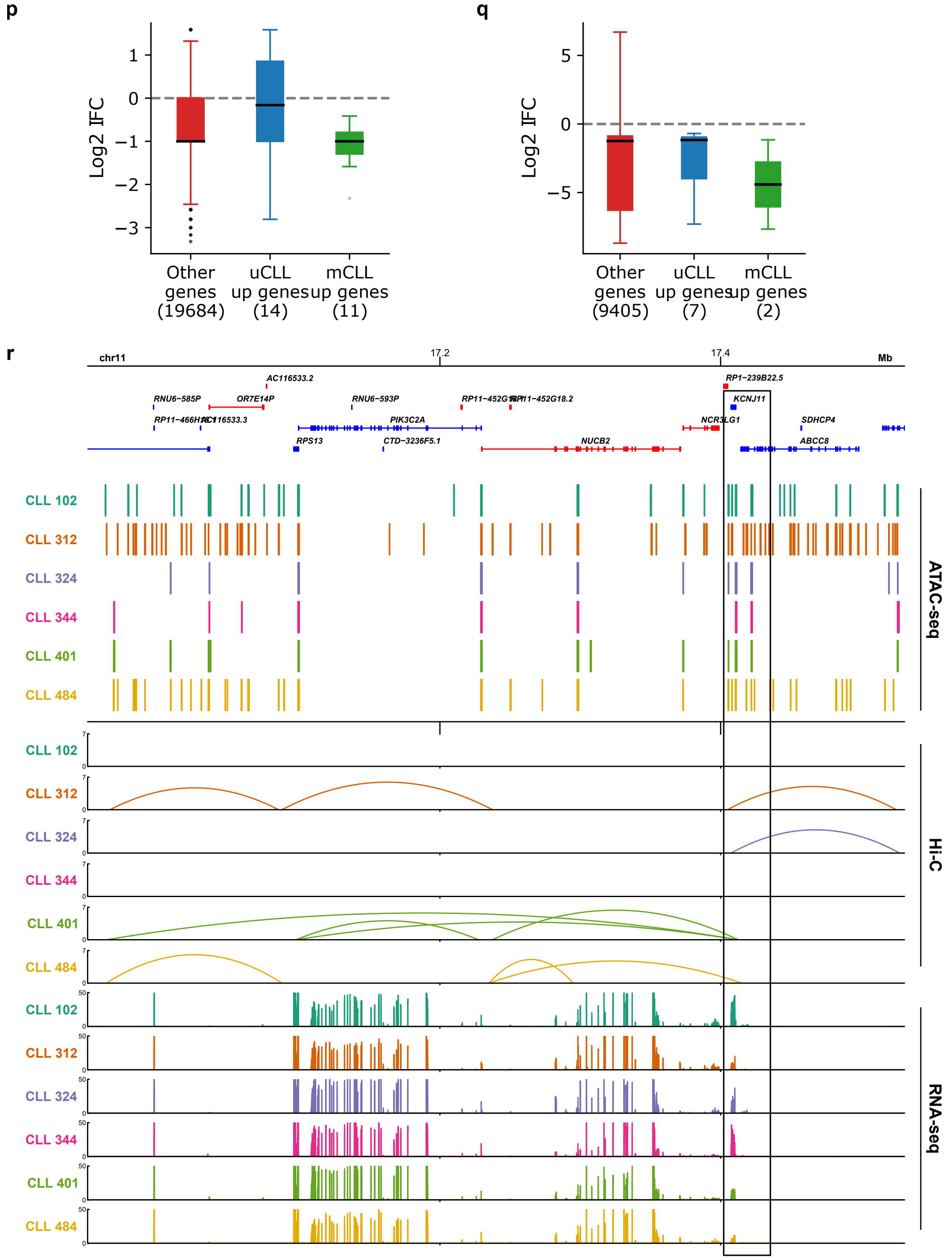

Supplementary Figure 8 p-r

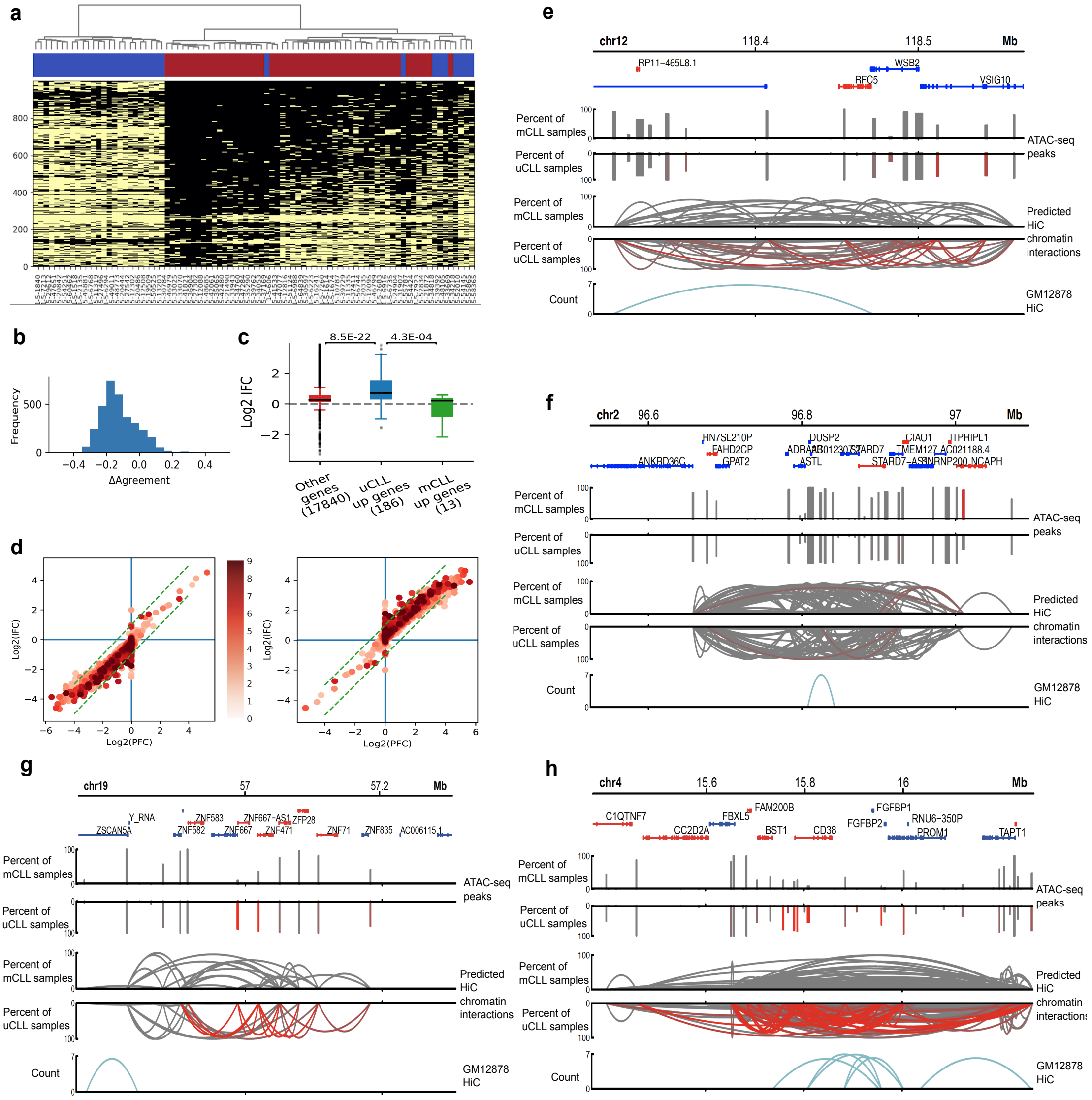

Supplementary Figure 9

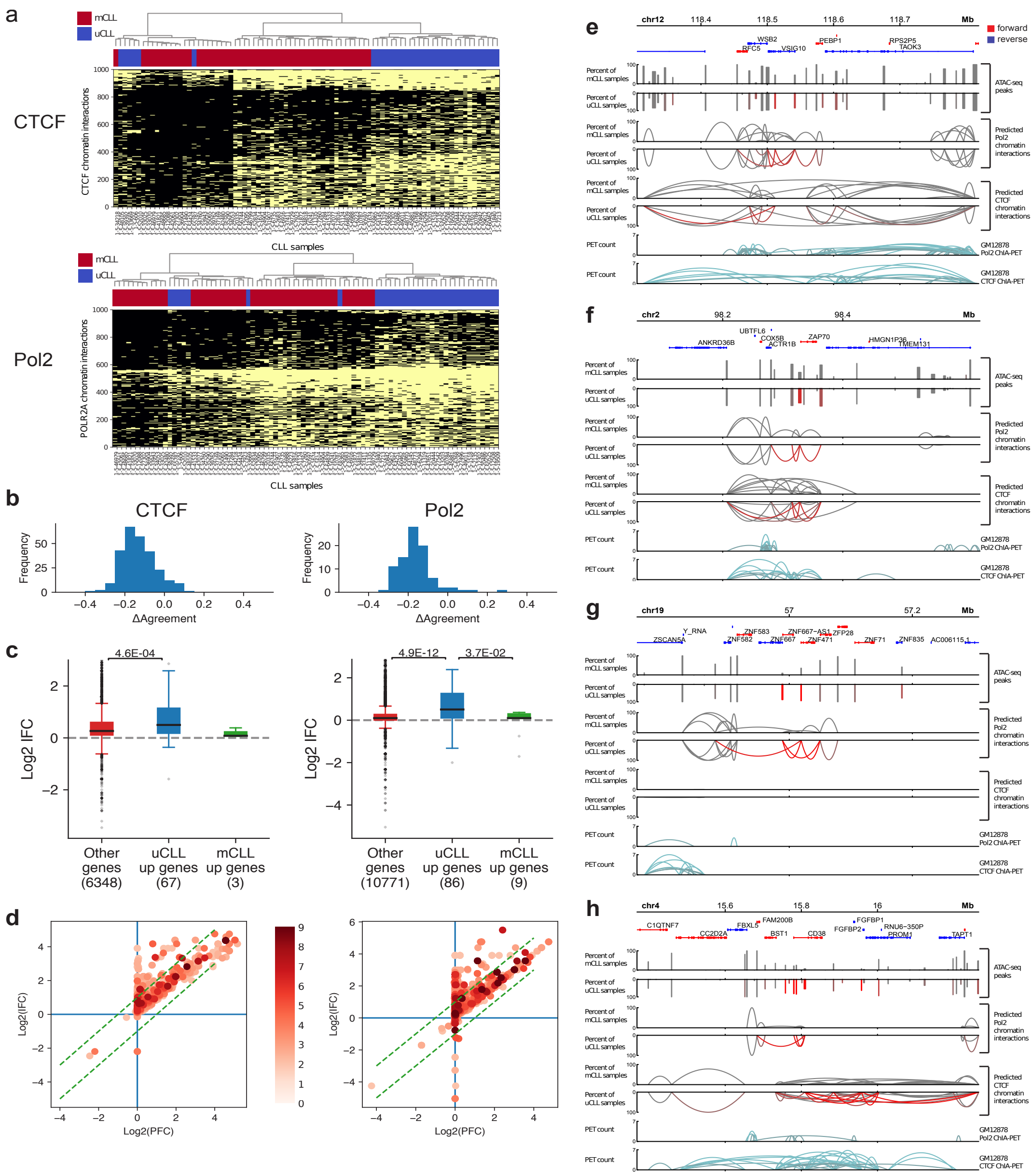

Supplementary Figure 10

#### 5 Supplementary Table

### Supplementary Table 1

| GM12878 | K562 | Helas3 |
|---|---|---|
| AwgTfbsHaibGm12878Cebpbsc150V0422111UniPk | AwgTfbsHaibK562Cebpbsc150V0422111UniPk | AwgTfbsSydhHelas3CebpbIggrabUniPk |
| AwgTfbsSydhGm12878CfosUniPk | AwgTfbsSydhK562CfosUniPk | AwgTfbsSydhHelas3CfosUniPk |
| AwgTfbsSydhGm12878Chd2ab68301IggmusUniPk | AwgTfbsSydhK562Chd2ab68301IggrabUniPk | AwgTfbsSydhHelas3Chd2IggrabUniPk |
| AwgTfbsUtaGm12878CmycUniPk | AwgTfbsUtaK562CmycUniPk | AwgTfbsUtaHelas3CmycUniPk |
| AwgTfbsSydhGm12878Corestsc30189IggmusUniPk | AwgTfbsSydhK562Corestsc30189IggrabUniPk | AwgTfbsSydhHelas3Corestsc30189IggrabUniPk |
| AwgTfbsBroadGm12878CtcfUniPk | AwgTfbsBroadK562CtcfUniPk | AwgTfbsBroadHelas3CtcfUniPk |
| AwgTfbsSydhGm12878E2f4IggmusUniPk | AwgTfbsSydhK562E2f4UcdUniPk | AwgTfbsSydhHelas3E2f4UniPk |
| AwgTfbsSydhGm12878Elk112771IggmusUniPk | AwgTfbsSydhK562Elk112771IggrabUniPk | AwgTfbsSydhHelas3Elk112771IggrabUniPk |
| AwgTfbsBroadGm12878Ezh239875UniPk | AwgTfbsBroadK562Ezh239875UniPk | AwgTfbsBroadHelas3Ezh239875UniPk |
| AwgTfbsHaibGm12878GabpPcr2xUniPk | AwgTfbsHaibK562GabpV0416101UniPk | AwgTfbsHaibHelas3GabpPcr1xUniPk |
| BroadHistoneGm12878H3k27acStdAl | BroadHistoneK562H3k27acStdAl | BroadHistoneHelas3H3k27acStdAl |
| BroadHistoneGm12878H3k27me3StdAl | BroadHistoneK562H3k27me3StdAl | BroadHistoneHelas3H3k27me3StdAl |
| BroadHistoneGm12878H3k36me3StdAl | BroadHistoneK562H3k36me3StdAl | BroadHistoneHelas3H3k36me3StdAl |
| BroadHistoneGm12878H3k4me2StdAl | BroadHistoneK562H3k4me2StdAl | BroadHistoneHelas3H3k4me2StdAl |
| BroadHistoneGm12878H3k4me3StdAl | BroadHistoneK562H3k4me3StdAl | BroadHistoneHelas3H3k4me3StdAl |
| BroadHistoneGm12878H3k79me2StdAl | BroadHistoneK562H3k79me2StdAl | BroadHistoneHelas3H3k79me2StdAl |
| BroadHistoneGm12878H3k9acStdAl | BroadHistoneK562H3k9acStdAl | BroadHistoneHelas3H3k9acStdAl |
| BroadHistoneGm12878H4k20me1StdAl | BroadHistoneK562H4k20me1StdAl | BroadHistoneHelas3H4k20me1StdAl |
| AwgTfbsSydhGm12878JundUniPk | AwgTfbsSydhK562JundIggrabUniPk | AwgTfbsSydhHelas3JundIggrabUniPk |
| AwgTfbsSydhGm12878MaxIggmusUniPk | AwgTfbsSydhK562MaxIggrabUniPk | AwgTfbsSydhHelas3MaxIggrabUniPk |
| AwgTfbsSydhGm12878Mazab85725IggmusUniPk | AwgTfbsSydhK562Mazab85725IggrabUniPk | AwgTfbsSydhHelas3Mazab85725IggrabUniPk |
| AwgTfbsSydhGm12878Mxi1IggmusUniPk | AwgTfbsSydhK562Mxi1af4185IggrabUniPk | AwgTfbsSydhHelas3Mxi1af4185IggrabUniPk |
| AwgTfbsSydhGm12878NfyaIggmusUniPk | AwgTfbsSydhK562NfyaUniPk | AwgTfbsSydhHelas3NfyaIggrabUniPk |
| AwgTfbsSydhGm12878NfybIggmusUniPk | AwgTfbsSydhK562NfybUniPk | AwgTfbsSydhHelas3NfybIggrabUniPk |
| AwgTfbsSydhGm12878Nrf1IggmusUniPk | AwgTfbsSydhK562Nrf1IggrabUniPk | AwgTfbsSydhHelas3Nrf1IggmusUniPk |
| AwgTfbsHaibGm12878NrsfPcr1xUniPk | AwgTfbsHaibK562NrsfV0416102UniPk | AwgTfbsHaibHelas3NrsfPcr1xUniPk |
| AwgTfbsSydhGm12878P300IggmusUniPk | AwgTfbsSydhK562P300IggrabUniPk | AwgTfbsSydhHelas3P300sc584sc584IggrabUniPk |
| AwgTfbsSydhGm12878Pol2UniPk | AwgTfbsSydhK562Pol2UniPk | AwgTfbsSydhHelas3Pol2UniPk |
| AwgTfbsSydhGm12878Pol2s2IggmusUniPk | AwgTfbsSydhK562Pol2s2IggrabUniPk | AwgTfbsSydhHelas3Pol2s2IggrabUniPk |
| AwgTfbsSydhGm12878Rad21IggrabUniPk | AwgTfbsSydhK562Rad21UniPk | AwgTfbsSydhHelas3Rad21IggrabUniPk |
| AwgTfbsSydhGm12878Rfx5200401194IggmusUniPk | AwgTfbsSydhK562Rfx5IggrabUniPk | AwgTfbsSydhHelas3Rfx5200401194IggrabUniPk |
| AwgTfbsSydhGm12878Smc3ab9263IggmusUniPk | AwgTfbsSydhK562Smc3ab9263IggrabUniPk | AwgTfbsSydhHelas3Smc3ab9263IggrabUniPk |
| AwgTfbsHaibGm12878Taf1Pcr1xUniPk | AwgTfbsHaibK562Taf1V0416101UniPk | AwgTfbsHaibHelas3Taf1Pcr1xUniPk |
| AwgTfbsSydhGm12878TbpIggmusUniPk | AwgTfbsSydhK562TbpIggmusUniPk | AwgTfbsSydhHelas3TbpIggrabUniPk |
| AwgTfbsSydhGm12878Tr4UniPk | AwgTfbsSydhK562Tr4UcdUniPk | AwgTfbsSydhHelas3Tr4UniPk |
| AwgDnaseUwdukeGm12878UniPk | AwgDnaseUwdukeK562UniPk | AwgDnaseUwdukeHelas3UniPk |
| AwgTfbsSydhGm12878Usf2IggmusUniPk | AwgTfbsSydhK562Usf2IggrabUniPk | AwgTfbsSydhHelas3Usf2IggmusUniPk |
| AwgTfbsSydhGm12878Znf143166181apUniPk | AwgTfbsSydhK562Znf143IggrabUniPk | AwgTfbsSydhHelas3Znf143IggrabUniPk |
| AwgTfbsSydhGm12878Znf274UniPk | AwgTfbsSydhK562Znf274UcdUniPk | AwgTfbsSydhHelas3Znf274UcdUniPk |

Supplementary Table 2

| | GM12878 | HeLaS3 | | HMEC | HUVEC | | IMR90 | K562 | | KBM7 | | NHEK |
|---|---|---|---|---|---|---|---|---|---|---|---|---|
| motif | CTCF | CTCF | MYOG | CTCF | CTCF | HES7 | CTCF | CTCF | MXI1 | CTCF | RXRB | CTCF |
| | CTCFL | CTCFL | HEN1 | CTCFL | CTCFL | SP4 | CTCFL | CTCFL | MYC | CTCFL | NR1H2 | CTCFL |
| | FOS | FOSL1 | ITF2 | JUNB | JUNB | KLF14 | JUNB | FOS | HEY1 | FOS | ERR3 | JUNB |
| | JUNB | FOS | ESR2 | FOS | FOS | TFE2 | FOS | JUNB | HES7 | JUNB | RARG | FOS |
| | ZN436 | JUNB | HMX3 | JUND | FOSL1 | ITF2 | JUND | FOSL1 | BMAL1 | JUN | RXRG | JUND |
| | JUN | JUN | MYF6 | FOSL1 | JUN | HEN1 | ZN436 | JUN | HES5 | FOSL1 | MAX | JUN |
| | JUND | JUND | ZBT18 | JUN | JUND | MYOG | JUN | JUND | HES1 | JUND | HES1 | FOSL1 |
| | FOSL1 | FOSL2 | KLF13 | FOSL2 | ZN436 | MYF6 | FOSB | ZN436 | RXRG | ZN436 | NR6A1 | ZN436 |
| | FOSL2 | ZN436 | MAX | FOSB | FOSL2 | HES1 | FOSL1 | FOSL2 | MAX | FOSL2 | ZBT18 | FOSB |
| | FOSB | FOSB | PTF1A | BACH2 | FOSB | ZBT18 | BACH2 | FOSB | SP4 | FOSB | SP4 | FOSL2 |
| | NFE2 | NF2L2 | RARA | BACH1 | INSM1 | MAX | NFE2 | NF2L2 | KLF14 | NF2L2 | CLOCK | NFE2 |
| | BACH2 | CR3L1 | NR6A1 | NF2L2 | NFE2 | | MAFK | NFE2 | PTF1A | NFE2 | KLF14 | BACH2 |
| | NF2L2 | NFE2 | HMX2 | NFE2 | NF2L2 | | INSM1 | BACH2 | ESR2 | BACH2 | SP2 | NF2L2 |
| | MAFK | BACH2 | LBX2 | MAFK | BACH2 | | FOSL2 | INSM1 | CLOCK | INSM1 | ESR2 | INSM1 |
| | BACH1 | INSM1 | HES1 | INSM1 | MAFK | | BACH1 | MAFK | | MAFK | TYY1 | MAFK |
| | INSM1 | MAFK | KLF14 | HES7 | BACH1 | | NF2L2 | BACH1 | | BACH1 | | BACH1 |
| | TFEB | BACH1 | SP2 | HES5 | CR3L1 | | HEY2 | CR3L1 | | CR3L1 | | MAFG |
| | TFE3 | TFEB | SP4 | MYC | TFEB | | MYCN | TFEB | | TFEB | | TFEB |
| | USF1 | USF1 | ZN331 | HEY1 | USF1 | | CR3L1 | USF1 | | USF1 | | TFE3 |
| | USF2 | TFE3 | BHA15 | MITF | USF2 | | HES1 | TFE3 | | USF2 | | USF2 |
| | CR3L1 | USF2 | CLOCK | MAX | TFE3 | | MAFG | USF2 | | TFE3 | | USF1 |
| | MAFG | ERR2 | TYY1 | BMAL1 | MAFG | | ZBTB4 | MAFG | | MAFG | | CR3L1 |
| | HEY2 | NR4A2 | ESR1 | MYCN | KLF13 | | TFEB | MITF | | MITF | | HEY2 |
| | MYCN | NR4A3 | ASCL2 | USF1 | MITF | | TFE3 | KLF13 | | BATF | | MYCN |
| | KLF13 | THB | BHE40 | TFE3 | BATF | | USF2 | MYCN | | KLF13 | | MITF |
| | MITF | ERR1 | XBP1 | BHE40 | MYCN | | USF1 | HEY2 | | MYCN | | BATF |
| | TFAP4 | NR1H3 | SP1 | MXI1 | ERR2 | | | BATF | | ERR2 | | MYC |
| | HTF4 | MAFG | KLF9 | CR3L1 | NR4A2 | | | TFAP4 | | NR4A2 | | MXI1 |
| | MYOD1 | MYCN | TFDP1 | RFX3 | THB | | | MYOD1 | | THB | | MAX |
| | BATF | MITF | NR5A2 | MLXPL | NR4A3 | | | HTF4 | | NR1H3 | | HES1 |
| | TFE2 | BATF | TGIF2 | USF2 | NR1H3 | | | ERR2 | | NR4A3 | | NR4A2 |
| | ITF2 | THA | ZIC3 | SNAI2 | ERR1 | | | NR4A2 | | HEY2 | | THB |
| | SP4 | NR1H2 | COT2 | HEY2 | TFAP4 | | | THB | | ERR1 | | ERR2 |
| | SP2 | PPARA | PAX3 | TFEB | MYOD1 | | | NR1H3 | | MXI1 | | |
| | KLF14 | RXRB | PAX7 | MLX | HTF4 | | | NR4A3 | | HEY1 | | |
| | MXI1 | ERR3 | HXA1 | BATF | HEY2 | | | ERR1 | | BMAL1 | | |
| | HES1 | RXRG | | SNAI1 | PPARA | | | TFE2 | | MYC | | |
| | MYC | RARG | | CLOCK | THA | | | HEN1 | | HES5 | | |
| | HEY1 | RXRA | | HES1 | RXRB | | | ITF2 | | HES7 | | |
| | BMAL1 | HEY1 | | BHE41 | RXRA | | | MYOG | | TFAP4 | | |
| | HES5 | HEY2 | | MAFG | ERR3 | | | MYF6 | | MYOD1 | | |
| | HES7 | HES5 | | | NR1H2 | | | THA | | HTF4 | | |
| | MAX | HES7 | | | RXRG | | | PPARA | | TFE2 | | |
| | MYOG | BMAL1 | | | NR6A1 | | | RXRB | | ITF2 | | |
| | HEN1 | MYC | | | RARG | | | RXRA | | HEN1 | | |
| | MYF6 | MXI1 | | | MXI1 | | | ERR3 | | MYOG | | |
| | ZBT18 | TFAP4 | | | HEY1 | | | NR1H2 | | MYF6 | | |
| | CLOCK | MYOD1 | | | BMAL1 | | | NR6A1 | | PPARA | | |
| | SP1 | HTF4 | | | MYC | | | RARG | | THA | | |
| | TFDP1 | TFE2 | | | HES5 | | | ZBT18 | | RXRA | | |

### Supplementary Table 3

| Motif | Number of samples with the motif detected | GM12878 | HeLaS3 | HMEC | HUVEC | IMR90 | K562 | KBM7 | NHEK |
|---|---|---|---|---|---|---|---|---|---|
| CTCF | 8 | 2.69E-06 | 1.08E-05 | 5.40E-08 | 4.41E-07 | 2.88E-08 | 8.27E-07 | 1.75E-06 | 7.02E-08 |
| CTCFL | 8 | 2.90E-06 | 1.08E-05 | 2.42E-05 | 1.53E-05 | 2.65E-05 | 9.93E-06 | 1.07E-05 | 1.46E-05 |
| FOS | 8 | 2.12E-04 | 1.30E-04 | 5.16E-04 | 1.60E-04 | 4.42E-04 | 1.80E-04 | 1.75E-04 | 1.24E-04 |
| JUNB | 8 | 2.12E-04 | 1.30E-04 | 5.16E-04 | 1.60E-04 | 4.42E-04 | 1.80E-04 | 1.75E-04 | 1.24E-04 |
| JUN | 8 | 3.38E-04 | 1.30E-04 | 6.09E-04 | 1.68E-04 | 1.85E-03 | 1.80E-04 | 1.86E-04 | 4.03E-04 |
| JUND | 8 | 3.38E-04 | 1.30E-04 | 5.81E-04 | 1.68E-04 | 1.61E-03 | 1.80E-04 | 1.86E-04 | 3.54E-04 |
| FOSL1 | 8 | 3.85E-04 | 1.30E-04 | 6.09E-04 | 1.68E-04 | 2.19E-03 | 1.80E-04 | 1.86E-04 | 4.03E-04 |
| FOSL2 | 8 | 7.08E-04 | 1.72E-04 | 8.17E-04 | 4.26E-04 | 3.85E-03 | 4.03E-04 | 3.62E-04 | 9.05E-04 |
| FOSB | 8 | 7.08E-04 | 5.56E-04 | 8.17E-04 | 5.47E-04 | 1.85E-03 | 7.32E-04 | 6.44E-04 | 8.09E-04 |
| NFE2 | 8 | 1.11E-03 | 2.43E-03 | 1.83E-03 | 1.13E-03 | 2.22E-03 | 1.15E-03 | 1.27E-03 | 1.15E-03 |
| BACH2 | 8 | 1.63E-03 | 2.43E-03 | 1.33E-03 | 1.13E-03 | 2.22E-03 | 1.20E-03 | 1.31E-03 | 1.15E-03 |
| NF2L2 | 8 | 2.01E-03 | 1.07E-03 | 1.83E-03 | 1.13E-03 | 6.13E-03 | 1.08E-03 | 9.61E-04 | 1.58E-03 |
| MAFK | 8 | 2.68E-03 | 7.99E-03 | 1.83E-03 | 3.40E-03 | 2.52E-03 | 3.97E-03 | 4.30E-03 | 2.40E-03 |
| BACH1 | 8 | 3.34E-03 | 8.70E-03 | 1.83E-03 | 4.59E-03 | 4.15E-03 | 5.35E-03 | 5.24E-03 | 4.11E-03 |
| INSM1 | 8 | 4.16E-03 | 2.73E-03 | 8.78E-03 | 1.01E-03 | 3.24E-03 | 1.51E-03 | 1.69E-03 | 1.96E-03 |
| TFEB | 8 | 1.30E-02 | 1.26E-02 | 3.39E-02 | 9.49E-03 | 4.56E-02 | 9.72E-03 | 9.10E-03 | 1.54E-02 |
| TFE3 | 8 | 1.30E-02 | 1.26E-02 | 1.22E-02 | 9.49E-03 | 4.56E-02 | 9.72E-03 | 9.10E-03 | 1.54E-02 |
| USF1 | 8 | 1.30E-02 | 1.26E-02 | 1.22E-02 | 9.49E-03 | 4.97E-02 | 9.72E-03 | 9.10E-03 | 1.54E-02 |
| USF2 | 8 | 1.30E-02 | 1.26E-02 | 3.39E-02 | 9.49E-03 | 4.97E-02 | 9.72E-03 | 9.10E-03 | 1.54E-02 |
| CR3L1 | 8 | 1.55E-02 | 2.11E-03 | 1.72E-02 | 7.98E-03 | 4.29E-02 | 8.44E-03 | 6.61E-03 | 1.77E-02 |
| MAFG | 8 | 1.74E-02 | 1.91E-02 | 3.94E-02 | 1.44E-02 | 4.33E-02 | 1.27E-02 | 1.65E-02 | 1.34E-02 |
| HEY2 | 8 | 2.57E-02 | 2.08E-02 | 3.39E-02 | 3.99E-02 | 4.29E-02 | 2.86E-02 | 3.04E-02 | 2.94E-02 |
| MYCN | 8 | 2.57E-02 | 1.99E-02 | 1.22E-02 | 3.48E-02 | 4.29E-02 | 2.86E-02 | 2.82E-02 | 2.94E-02 |
| HES1 | 8 | 4.10E-02 | 3.67E-02 | 3.62E-02 | 4.64E-02 | 4.29E-02 | 3.89E-02 | 3.79E-02 | 4.86E-02 |
| ZN436 | 7 | 3.14E-04 | 2.77E-04 | - | 3.85E-04 | 1.76E-03 | 2.51E-04 | 2.82E-04 | 5.38E-04 |
| MITF | 7 | 3.18E-02 | 1.99E-02 | 8.85E-03 | 2.59E-02 | - | 2.41E-02 | 2.48E-02 | 3.45E-02 |
| BATF | 7 | 3.85E-02 | 2.01E-02 | 3.43E-02 | 2.77E-02 | - | 2.88E-02 | 2.59E-02 | 3.86E-02 |
| MXI1 | 7 | 4.10E-02 | 2.08E-02 | 1.72E-02 | 4.32E-02 | - | 3.89E-02 | 3.34E-02 | 4.84E-02 |
| MYC | 7 | 4.10E-02 | 2.08E-02 | 8.85E-03 | 4.32E-02 | - | 3.89E-02 | 3.34E-02 | 4.84E-02 |
| MAX | 7 | 4.10E-02 | 2.75E-02 | 1.21E-02 | 4.81E-02 | - | 4.22E-02 | 3.79E-02 | 4.84E-02 |
| HEY1 | 6 | 4.10E-02 | 2.08E-02 | 8.85E-03 | 4.32E-02 | - | 3.89E-02 | 3.34E-02 | - |
| BMAL1 | 6 | 4.10E-02 | 2.08E-02 | 1.21E-02 | 4.32E-02 | - | 3.89E-02 | 3.34E-02 | - |
| HES5 | 6 | 4.10E-02 | 2.08E-02 | 8.85E-03 | 4.32E-02 | - | 3.89E-02 | 3.34E-02 | - |
| HES7 | 6 | 4.10E-02 | 2.08E-02 | 8.85E-03 | 4.32E-02 | - | 3.89E-02 | 3.34E-02 | - |
| KLF13 | 5 | 2.83E-02 | 2.74E-02 | - | 2.51E-02 | - | 2.74E-02 | 2.76E-02 | - |
| TFAP4 | 5 | 3.36E-02 | 2.35E-02 | - | 3.92E-02 | - | 3.29E-02 | 3.49E-02 | - |
| HTF4 | 5 | 3.36E-02 | 2.35E-02 | - | 3.92E-02 | - | 3.29E-02 | 3.49E-02 | - |
| MYOD1 | 5 | 3.36E-02 | 2.35E-02 | - | 3.92E-02 | - | 3.29E-02 | 3.49E-02 | - |
| TFE2 | 5 | 3.87E-02 | 2.36E-02 | - | 4.52E-02 | - | 3.45E-02 | 3.68E-02 | - |
| ITF2 | 5 | 3.89E-02 | 2.36E-02 | - | 4.52E-02 | - | 3.45E-02 | 3.68E-02 | - |
| SP4 | 5 | 3.91E-02 | 3.74E-02 | - | 4.46E-02 | - | 4.41E-02 | 4.64E-02 | - |
| KLF14 | 5 | 3.91E-02 | 3.74E-02 | - | 4.46E-02 | - | 4.41E-02 | 4.70E-02 | - |
| MYOG | 5 | 4.46E-02 | 2.36E-02 | - | 4.52E-02 | - | 3.45E-02 | 3.68E-02 | - |
| HEN1 | 5 | 4.46E-02 | 2.36E-02 | - | 4.52E-02 | - | 3.45E-02 | 3.68E-02 | - |
| MYF6 | 5 | 4.46E-02 | 2.64E-02 | - | 4.52E-02 | - | 3.45E-02 | 3.68E-02 | - |
| ZBT18 | 5 | 4.46E-02 | 2.64E-02 | - | 4.71E-02 | - | 3.64E-02 | 4.19E-02 | - |
| CLOCK | 5 | 4.62E-02 | 3.81E-02 | 3.53E-02 | - | - | 4.96E-02 | 4.65E-02 | - |
| ERR2 | 5 | - | 1.55E-02 | - | 3.68E-02 | - | 3.30E-02 | 2.99E-02 | 4.89E-02 |
| NR4A2 | 5 | - | 1.55E-02 | - | 3.68E-02 | - | 3.30E-02 | 2.99E-02 | 4.89E-02 |
| THB | 5 | - | 1.55E-02 | - | 3.68E-02 | - | 3.45E-02 | 2.99E-02 | 4.89E-02 |
| NR4A3 | 4 | - | 1.55E-02 | - | 3.68E-02 | - | 3.45E-02 | 2.99E-02 | - |
| ERR1 | 4 | - | 1.55E-02 | - | 3.68E-02 | - | 3.45E-02 | 3.18E-02 | - |
| NR1H3 | 4 | - | 1.58E-02 | - | 3.68E-02 | - | 3.45E-02 | 2.99E-02 | - |
| THA | 4 | - | 2.02E-02 | - | 4.08E-02 | - | 3.62E-02 | 3.73E-02 | - |
| NR1H2 | 4 | - | 2.02E-02 | - | 4.08E-02 | - | 3.62E-02 | 3.73E-02 | - |
| PPARA | 4 | - | 2.02E-02 | - | 4.08E-02 | - | 3.62E-02 | 3.73E-02 | - |
| RXRB | 4 | - | 2.02E-02 | - | 4.08E-02 | - | 3.62E-02 | 3.73E-02 | - |
| ERR3 | 4 | - | 2.02E-02 | - | 4.08E-02 | - | 3.62E-02 | 3.73E-02 | - |
| RXRG | 4 | - | 2.02E-02 | - | 4.08E-02 | - | 4.15E-02 | 3.73E-02 | - |
| RARG | 4 | - | 2.02E-02 | - | 4.08E-02 | - | 3.62E-02 | 3.73E-02 | - |

| | | | | | | | | | |
|---|---|---|---|---|---|---|---|---|---|
| RXRA | 4 | - | 2.02E-02 | - | 4.08E-02 | - | 3.62E-02 | 3.73E-02 | - |
| NR6A1 | 4 | - | 3.18E-02 | - | 4.08E-02 | - | 3.62E-02 | 4.10E-02 | - |
| SP2 | 3 | 3.91E-02 | 3.74E-02 | - | - | - | - | 4.70E-02 | - |
| ESR2 | 3 | - | 2.40E-02 | - | - | - | 4.68E-02 | 4.74E-02 | - |
| SP1 | 2 | 4.64E-02 | 4.23E-02 | - | - | - | - | - | - |
| TFDP1 | 2 | 4.64E-02 | 4.44E-02 | - | - | - | - | - | - |
| PTF1A | 2 | - | 3.09E-02 | - | - | - | 4.62E-02 | - | - |
| TYY1 | 2 | - | 3.90E-02 | - | - | - | - | 4.93E-02 | - |
| BHE40 | 2 | - | 4.03E-02 | 1.72E-02 | - | - | - | - | - |
| HMX3 | 1 | - | 2.57E-02 | - | - | - | - | - | - |
| RARA | 1 | - | 3.18E-02 | - | - | - | - | - | - |
| HMX2 | 1 | - | 3.58E-02 | - | - | - | - | - | - |
| LBX2 | 1 | - | 3.58E-02 | - | - | - | - | - | - |
| ZN331 | 1 | - | 3.80E-02 | - | - | - | - | - | - |
| BHA15 | 1 | - | 3.80E-02 | - | - | - | - | - | - |
| ESR1 | 1 | - | 3.92E-02 | - | - | - | - | - | - |
| ASCL2 | 1 | - | 4.02E-02 | - | - | - | - | - | - |
| XBP1 | 1 | - | 4.04E-02 | - | - | - | - | - | - |
| KLF9 | 1 | - | 4.23E-02 | - | - | - | - | - | - |
| NR5A2 | 1 | - | 4.49E-02 | - | - | - | - | - | - |
| TGIF2 | 1 | - | 4.54E-02 | - | - | - | - | - | - |
| ZIC3 | 1 | - | 4.54E-02 | - | - | - | - | - | - |
| COT2 | 1 | - | 4.64E-02 | - | - | - | - | - | - |
| PAX3 | 1 | - | 4.83E-02 | - | - | - | - | - | - |
| PAX7 | 1 | - | 4.83E-02 | - | - | - | - | - | - |
| HXA1 | 1 | - | 4.83E-02 | - | - | - | - | - | - |
| RFX3 | 1 | - | - | 2.46E-02 | - | - | - | - | - |
| MLXPL | 1 | - | - | 2.87E-02 | - | - | - | - | - |
| SNAI2 | 1 | - | - | 3.39E-02 | - | - | - | - | - |
| MLX | 1 | - | - | 3.39E-02 | - | - | - | - | - |
| SNAI1 | 1 | - | - | 3.53E-02 | - | - | - | - | - |
| BHE41 | 1 | - | - | 3.62E-02 | - | - | - | - | - |
| ZBTB4 | 1 | - | - | - | - | 4.51E-02 | - | - | - |

Supplementary Table 4

| Sample | Percentage of similarity to *IGHV* reference sequence |
|---|---|
| AD102 | 98.61% |
| AD312 | 93.06% |
| AD324 | 97.22% |
| AD344 | 100.00% |
| AD401 | 64.91% |
| AD484 | 90.28% |

Supplementary Table 5

| | Sample | 102 | 312 | 324 | 344 | 401 | 484 |
|---|---|---|---|---|---|---|---|
| Read alignment statistics | Total reads | 948,029,258 | 952,568,936 | 962,597,696 | 928,396,764 | 921,586,154 | 943,118,124 |
| | Normal paired | 230,671,868 | 229,040,734 | 227,975,418 | 190,027,764 | 241,214,740 | 226,157,764 |
| | Ligation motif present | 535,901,842 | 479,621,330 | 493,454,930 | 515,211,736 | 480,234,368 | 509,891,762 |
| | Alignable | 858,223,558 | 867,092,356 | 883,427,000 | 844,824,800 | 832,474,774 | 854,272,762 |
| Hi-C analysis statistics by Juicer | Unique reads | 348,873,216 | 367,167,312 | 376,033,377 | 347,467,945 | 349,407,755 | 357,147,702 |
| | Hi-C contacts | 314,771,996 | 330,897,673 | 341,339,626 | 314,806,260 | 314,697,476 | 321,307,304 |
| | Inter-chromosomal | 72,179,180 | 85,636,523 | 85,782,728 | 62,185,458 | 59,258,481 | 64,967,149 |
| | Intra-chromosomal | 242,592,816 | 245,261,150 | 255,556,898 | 252,620,802 | 255,438,995 | 256,340,155 |
| | Short range (<20kb) | 110,379,302 | 92,854,062 | 110,974,633 | 101,510,722 | 112,777,329 | 120,382,809 |
| | Long range(>20kb) | 132,212,196 | 152,406,643 | 144,581,318 | 151,109,799 | 142,660,579 | 135,956,351 |
| | # contact domains (10k) | 1,339 | 1,133 | 1,591 | 1,649 | 1,796 | 1,583 |
| | # loops identified | 8,329 | 5,156 | 6,981 | 8,197 | 12,710 | 12,700 |

Supplementary Table 6

| IGHV mutation test primers | | |
|---|---|---|
| 5’ IGHV leader primers | IGHV1L | AAATCGATACCACCATGGACTGGACCTGGAGG |
| | IGHV1bL | AAATCGATACCACCATGGACTGGACCTGGAG(C/A) |
| | IGHV2aL | AAATCGATACCACCATGGACACACTTTGCT(A/C)AC |
| | IGHV2bL | AAATCGATACCACCATGGACATACTTTGTTCCAC |
| | IGHV3aL | AAATCGATACCACCACCATGGAGTTTGGGCTGAGC |
| | IGHV3bL | AAATCGATACCACCACCATGGA(A/G)(C/T)T(G/T)(G/T)G(G/A)CT(G/C/ 5 T)(A/C/T)GC |
| | IGHV4L | AAATCGATACCACCATGAAACACCTGTGGTTCTT |
| | IGHV5L | AAATCGATACCACCATGGGGTCAACCGCCATC |
| | IGHV6L | AAATCGATACCACCATGTCTGTCTCCTTCCTC |
| 3’ IGHJ primers | IGHJ1-2 | TGAGGAGACGGTGACCAGGGTGCC |
| | IGHJ3 | TGAAGAGACGGTGACCATTGTCCC |
| | IGHJ4-5 | TGAGGAGACGGTGACCAGGGTTCC |
| | IGHJ6 | TGAGGAGACGGTGACCGTGGTCCC |

| 4C viewpoints and primers in MCF-7 cells | | | |
|---|---|---|---|
| | CAP2 | ELOVL2 | GREB1 |
| Viewpoint | chr6:17390954-17391643 | chr6:11047037-11047317 | chr2:11691065-11691295 |
| Outer Forward | GGGCTGATAACACAGTAGACTCA | GTGCTTTGCAAGGTTTAGATGACAT | AGAGGACTCGGGTAGAATGAGAA |
| Outer Reverse | GACAAAGGAGATGACGGGAGAAA | ACTTCCACATTACCATTTAGCCTGA | ATAGCTCAGGTCAATGTCCCAG |
| Nested Forward | GAATCTCAACAGAGGAGGTGACC | TTCTTAGAGGTCGGTTGTCACTT | TCAACTAGAATAAGTGCAGGGC |
| Nested Reverse | CACAGTTATCAGCCTCCTTGTCT | TACACATCAACCCATAAGGCCAG | TGGGTCAGTTGGCTTCATCT |

| | SIAH2 | P2RY2 |
|---|---|---|
| Viewpoint | chr3:150478089-150478323 | chr11:72933550-72934990 |
| Outer Forward | GCCTTTCTGTACATGTCAGTGGTT | CCTTGTGAACTCAGACTCAGATT |
| Outer Reverse | AGGTTGATTTCAGTAGCCTGTTTC | TGCTCCTCCTCAAATCTTATCAC |
| Nested Forward | GCTATTACTTTGCAAGCTGA | GGAGCCTGAGGAATTCACTTGAA |
| Nested Reverse | CCTGTTTCACGTTAGGAAAGAA | ATTGGAACACTAGACCTTGAGTAAG |
